# Planar cell polarity is essential for the architectural patterning of the mammalian biliary tree

**DOI:** 10.1101/2023.11.20.567871

**Authors:** Michaela Raab, Ersi Christodoulou, Roopesh Krishnankutty, Nicholas T Younger, Konstantinos Gournopanos, Alexander von Kriegsheim, Scott H Waddell, Luke Boulter

## Abstract

In the developing liver, bipotent epithelial progenitor cells known as hepatoblasts undergo lineage segregation to form the two major epithelial cell types, hepatocytes that constitute the bulk of the liver parenchyma and biliary epithelial cells (cholangiocytes) which comprise the bile duct, a complex tubular network which is critical for normal liver function. Notch and TGFβ signalling promote the formation of a sheet of biliary epithelial cells, the ductal plate that organises into discontinuous tubular structures. How these structures elongate and connect to form a continuous duct remains undefined. Here, we show that the planar cell polarity protein, VANGL2 is expressed late in intrahepatic bile duct development and patterns the formation of cell-cell contacts between biliary cells. The patterning of these cell contacts regulates the normal polarisation of the actin cytoskeleton within biliary cells and loss of *Vangl2*-function results in the abnormal distribution of cortical actin remodelling resulting in the failure of bile duct formation. Planar cell polarity is a critical step in the post-specification sculpture of the bile duct and is essential for establishing normal tissue architecture.

## Introduction

Intrahepatic bile ducts form in vertebrates from a transient embryological structure known as the ductal plate that comprises two layers of simple epithelial cells^1^. The developmental signals that are required to specify the ductal plate from bipotent hepatoblasts (foetal epithelial progenitor cells in the liver) are well known and deficiencies in both Notch and TGFβ signalling in particular are associated with poorly developed, mis-branched or absent bile ducts^2–5^. Alagille patients who have congenital mutations in *JAGGED1* and less frequently in *NOTCH2,* for example, suffer from cholestasis and secondary liver disease due to a poorly formed biliary tree that necessitates non-curative surgery or liver transplantation^6^. Following specification of the biliary lineage, small stretches of primordial duct must lumenise to form discontinuous hollow tubes, which then elongate and intercalate to establish the final complex and branched biliary network^7^. What the molecular processes are that promote the formation of a continuous, higher-order ductular network from these discontinuous primordial ducts remains elusive.

Across a range of ductular or tubular tissues, including pancreas^8,9^, kidney^10^ and lung^11,12^, planar cell polarity (PCP) signalling is required for the collective polarisation and movement of epithelial cells. Loss or ectopic activation of PCP signalling is deleterious for normal tubular architecture, implying that cell-intrinsic levels of PCP components are critical for correct tissue patterning. In normal mammalian development, PCP proteins (including CELSR, VANGL and FZD, for example) asymmetrically localise along the proximal-distal axis of cells thereby imparting spatial information across a population of cells perpendicular to the apico-basal cell axis^13,14^. Indeed, evidence from zebrafish demonstrated that targeting PCP components *pk1a*, *vangl2* or *ankrd6* affects the development of a complex biliary tree^15^. While PCP confers a biochemical gradient across populations of cells within a tissue, how directionality is physically translated into polarised cellular movements is less clear. The prevailing hypothesis is that PCP proteins activate intracellular ROCK and RHO-GTPases^16^ to coordinate local cytoskeleton remodelling and cell-cell connectivity^17^.

We have previously demonstrated that following adult bile duct damage reactivation of PCP coordinates bile duct regrowth^18^, a process that recapitulates many of the features of bile duct ontogeny; therefore, we reasoned that during bile duct development, PCP could represent a critical factor in embryonic ductular patterning. Using a combination of single cell RNA sequencing data and a mutant mouse line carrying a hypomorphic mutation in Vangl2 (*Vangl2^S^*^464*N*^) we demonstrate that the expression of core PCP pathway components is restricted until late in development when the biliary tree is undergoing morphogenesis and by patterning cell-cell junctions, PCP drives terminal patterning of the bile duct.

## Materials and Methods

### Re-analysis of single cell data from Yang et al

TPM files were downloaded from GEO and analysed in R using the Seurat package. Prior to creating a Seurat object, duplicates were removed and cells were filtered. Cells with a unique feature count over 10,000 or less than 7,000 were removed. Following this, the data was normalised by applying the global-scaling normalisation method “LogNormalise()” which normalises the feature expression measurements for each cell by the total expression. The result is then multiplied by a scale factor of 10,000. Next, highly variable features were identified with the “FindVariableFeatures()” method, which returned 2000 features that exhibit high cell-to cell variation in the dataset and were used for downstream analysis. To determine whether cells cluster according to their cell cycle state, the function “CellCycleScoring” was applied, which revealed clustering of the cells based on their S– and G2M-Score. To overcome this, cell cycle regression was performed. Next, the linear transformation function “ScaleData” was applied to scale the data. This function shifts the expression of each gene so that the mean expression across the cells is 0 and scales the expression of each gene, such that the variance across cells is 1. For PCA analysis on the scaled data, the previously determined variable features were used as an input. To determine the dimensionality of the dataset, the “ElbowPlot()” was used, which ranks the principal components based on the percentage of variance. The elbow was found at around PC25-30, hence PC30 was chosen as a cut-off. For clustering the cells, the functions “FindNeighbours()” and “FindClusters()” were applied using previously defined dimensionality of the dataset (PC30) as input and at a resolution of 0.5. Non-linear dimensional reduction the UMAP technique was used, which identified 5 independent clusters. To find differentially expressed features, the function “FindAllMarkers() on positive markers was applied with a minimum percentage of 0.25 and a log fold change threshold of 0.25. To visualise marker expression, the functions “VlnPlot()”, “FeaturePlot()” and “DotPlot()” were used.

### Animal models

*Vangl2^+/GFP^* mice were kindly provided by Ping Chen and were maintained on a CD1 background^19^. *Vangl2^GFP/GFP^* (from hereon in called Vangl2^GFP^) embryos were used at E18.5 and *Vangl2^+/+^* littermates were used as controls*Vangl2^S464N^* mice: *Vangl2^+/S464N^* mice^20^ were provided by Harwell, UK and were maintained on a C3H background. Heterozygous animals were bred together to generate embryos homozygous for the *Vangl2* mutation, *Vangl2^S464N/S464N^* (abbreviated to *Vangl2^S464N^*) and *Vangl2^+/+^*controls were maintained on a C3H background. Embryonic days were counted from day of a found plug (E0.5) for both mouse lines.

Animals were maintained in SPF environment and studies carried out in accordance with the guidance issued by the Medical Research Council in “Responsibility in the Use of Animals in Medical Research” (July 1993) and licensed by the Home Office under the Animals (Scientific Procedures) Act 1986. Experiments were performed under project license number PFD31D3D4 in facilities at the University of Edinburgh (PEL 60/6025).

### Generation of Foetal Liver Organoids (FLO)

Livers were dissected from E15.5 *Vangl2^+/+^*, *Vangl2^S646N^* or *Vangl2^eGFP^*embryos under sterile conditions. Livers were digested with collagenase– and dispase-containing digestion buffer and dissociated into single cells. Pelleted cells were washed in PBS and suspended in 100% matrigel and added to a cell culture plate. Foetal liver cells were cultured in organoid culture media composed of Advanced DMEM/F-12 media supplemented with GlutaMAX, Antibiotic-Antimycotic, 10 μM HEPES, 50 ng/ml EGF, 100 ng/ml FGF10, 5 ng/ml HGF, 10 nM gastrin, 10 mM nicotinamide, 1.25 mM N-acetyl-L-cysteine, 1X B27, 1X N2 Supplement, 10 μM forskolin, 10 μM Y-27632, 5μM A83-01 and 3.33 μM Chir99021.

### Mass spectroscopy

Snap frozen E18.5 livers of *Vangl2^+/GFP^* and *Vangl2^+/+^* mice (N=3 in each case) were lysed in RIPA lysis buffer supplemented with protease inhibitor. Tissue was sonicated at 50Hz for 5min using a metal bead for each sample. The lysate was left at 4°C for 30 min to allow for complete cell lysis. Lysate was then centrifuged at 16.000 x g for 20 min at 4°C and protein lysate was transferred to a new tube. Protein concentration was measured by BCA assay. Co-immunoprecipitation pull down experiments (Co-IP) used 500 µl of protein lysate at a concentration of 2 mg/ml in lysis buffer. Co-IP was performed using the Kingsfisher Flex robot at the in-house mass spectrometry facility using the following protocol. To protein lysates, magnetic agarose GFP-Trap beads (Chromotek) were added to allow GFP binding to beads. Beads were washed in lysis buffer and protein eluted with TBS. Proteins were digested with trypsin and cysteine residues were alkylated with 2-Chloracetamide solution and kept in the dark. Columns for protein binding were prepared as follows: C18 Discs (Emmore 3M C18) were punched out using a blunted syringe needle and pushed into 200 μl tips before activating with methanol. Whole protein samples were loaded on to tip columns which were stored until mass spectrometry. Prior to mass spectrometry, protein was eluted from columns with 50% acetonitrile, dried and resuspended in 0.1 % TFA/Water and ran on a Lumos Fuison mass spectrometer coupled to a uHPLC RSLCnano (Thermo Fisher). To identify significantly enriched proteins, the median of the MaxLFQ.Intensity of each group (*Vangl2^eGFP^* and *Vangl2^+/+^*) between the three replicates was taken. The negative log fold change for both groups was calculated and subtracted from each other. Log fold change values >1 was considered as significantly enriched. All enriched proteins in the *Vangl2^eGFP^*samples were used for a downstream gene ontology analysis using the online DAVID and REVIGO platforms and compared against the *Vangl2^+/+^* control.

### Immunostaining of tissues and organoids

<u>FUnGi tissue clearing</u>^21^: E18.5 livers were dissected and either cut into thin slices or kept as whole livers and fixed in 4 % PFA. The tissue was transferred to Washing buffer (WB, PBS + 0.1 % Tween20 + 50 µg/ml ascorbic acid + 0.5 ng/ml reduced L-Glutathione) for 1 h at 4 °C rolling. Tissues were depigmented using DMSO, 30% H20 and PBS, (ratio of 1:2:4) and washed in Washing buffer 1 (WB1: PBS, 0.2 % Tween, 0.2 % Triton, 0.02 % SDS, 0.2 % BSA, 50 ug/ml ascorbic acid, 0.5 ng/ml L-glutathione reduced). Primary antibodies were diluted in washing buffer 2 (WB2: PBS + 0.1 % Triton-X-100 + 0.02 % SDS + 0.2 % BSA + 50 ug/ml ascorbic acid + 0.5 ng/ml L-Glutathione reduced) and incubated with tissue overnight. Livers were washed in WB2 and Secondary antibodies were diluted in WB2 (1:500) and incubated with livers ON at 4°C. Tissue was subsequently washed with WB2 and clarified with FunGI clearing agent (50/50 % v/v glycerol solution in H_2_O + 10.6 ml Tris Base + 1 mM EDTA + 2.5 M fructose + 2.5 M urea) overnight. Tissues were mounted on slides for imaging.

<u>IHC and DAB staining:</u> Dissected tissues were fixed overnight in formalin at 4 °C, embedded in paraffin and were sectioned at 4 µm. Following antigen retrieval (Supplementary Table 1), tissue sections were incubated with antibodies as detailed in Supplementary Table 1. Fluorescently stained tissues were counterstained with DAPI prior to imaging. Colorimetric stains were counterstained with haematoxylin and mounted with DPX. DAB mean measurements were quantified using QuPath (https://qupath.github.io/). Histological tissues were scanned using a Nanozoomer, using a Nikon A1R or Leica Stellaris confocal microscope and were analysed using either FIJI, Imaris, or QuPath.

### Organoid Immunofluorescent

Organoids were fixed with 4 % Formalin solution in glass-bottom plates. Following permeabilisation with Triton-X, cells were washed in PBS and glycine (PBS + 100 mM glycine) and proteins were blocked followed by incubation with primary antibodies, Supplementary Table 1. Organoids were mounted with Flouromount-G with DAPI prior to imaging.

### Electron Microscopy

Samples were fixed in 3% glutaraldehyde in 0.1 M Sodium Cacodylate buffer, pH 7.3, for 2 h then washed in three 10 min changes of 0.1 M Sodium Cacodylate. Specimens were then post-fixed in 1% Osmium Tetroxide in 0.1 M Sodium Cacodylate for 45 min, then washed in three 10 min changes of 0.1M Sodium Cacodylate buffer. These samples were then dehydrated in 50%, 70%, 90% and 100% ethanol (X3) for 15 min each, then in two 10 min changes in Propylene Oxide. Samples were then embedded in TAAB 812 resin. Sections, 1 μm thick were cut on a Leica Ultracut ultramicrotome, stained with Toluidine Blue, and viewed in a light microscope to select suitable areas for investigation. Ultrathin sections, 60nm thick were cut from selected areas, stained in Uranyl Acetate and Lead Citrate then viewed in a JEOL JEM-1400 Plus TEM. Representative images were collected on a GATAN OneView camera.

### Fluorescent cell membrane labelling

After establishing single cell suspensions of both *Vangl2^+/+^* and *Vangl2^S464N^* organoid lines, PKH26 (red) and PKH67 (green) general membrane dyes were used to label cells as per the manufacturer’s instructions. PKH67-labelled *Vangl2^+/+^* and PKH26-labelled *Vangl2^S464N^* cells were intermixed at a 1:1 ratio, with 20,000 cells in each well. Additionally, two control wells of PKH67-*Vangl2^+/+^*/PKH26-*Vangl2^+/+^* and PKH67-*Vangl2^S464N^*/PKH26-*Vangl2^S464N^* were also plated with the same cell density. After 2 days, images of formed organoids were acquired, and the number of red/green/mosaic organoids was recorded. A chi-square test was used to assess whether there were significantly meaningful differences between the three groups.

### Immunoblotting

Protein lysates were obtained from using RPPA lysis buffer (2.5 ml Triton-X-100, 25 ml 0.5 M HEPES pH 7.4, 0.5 ml 0.5 M EGTA pH 7.5-8.0, 37.5 ml 1 M sodium chloride, 0.375 ml 1M magnesium chloride, 0.1 ml 100 mM sodium orthovanadate, 1ml 100 mM tetrasodium pyrophosphate, 1 ml 1M sodium fluoride, 1 cOmplete mini EDTA-free protease inhibitor tablet (Roche), 1 phosphoSTOP phosphatase inhibitor tablet (Roche), 1 ml glycerol and 1.9 ml dH2O). For Western blots, lysates were loaded (7.5-20 µg protein) onto a 4-12% NuPAGE Bis-Tris gel (Thermo Fisher). Protein lysates were reduced with NuPAGE LDS sample buffer (4x) and NuPAGE Sample Reducing Agent (10x) prior to running. Gels were run using NuPAGE MOPS SDS Running buffer containing NuPAGE Antioxidant. Proteins were transferred onto PVDF membrane (Amersham) using NuPAGE Transfer buffer. Following transfer membranes were either blocked in 5% dried milk (Marvel) in PBST or 5% BSA (Sigma Aldrich). Membranes were incubated with primary antibodies (Supplementary Table 1) in 5% BSA (Sigma Aldrich) at 4 °C overnight. Following washing with PBST, membranes were incubated with HRP-conjugated secondary antibodies (Supplementary Table 1) in 3% dried milk (Marvel) or 3% BSA (Sigma Aldrich) at room temperature for 1 h. Following washing, signal was developed using ECL (Pierce) and visualised using Amersham ImageQuant 800 (Cytiva). Signal was quantified using either FIJI or Image Studio Lite (LI-COR).

### Live imaging of FLOs with/without SiR-actin

*Vangl2^+/+^* and *Vangl2^S464N^* organoids were dissociated into single cells and 5,000 cells for each FLO line were plated in organoid growth media glass bottom slide on a cushion of 1:1 Ultimatrix and PBS. 1 μM SiR-actin and 10 μM Verapamil was added to the organoid media. Organoids were imaged for 24 h. In assays where we assessed organoid growth, single-cell suspension was observed using the Incucyte S3 machine over a period of one week, and images were taken every 6 hours. Analysis was performed using the Incucyte S3 software.

## Results

### Planar Cell Polarity components are restricted to the ductular lineage in mammalian liver development

In the mouse, liver development is initiated from the foregut endoderm and following the formation of a liver bud at E10.5 liver epithelial cells undergo progressive specification and differentiation into the two principal epithelial cell lineage in the liver, hepatocytes and biliary epithelial cells (BECs, also known as cholangiocytes)^18,22,23^. Using a previously published data set in which epithelial cells were isolated using either DLK1 to select for hepatocellular lineages or EpCAM to enrich ductular cells (Figure 1A) we sought to determine the regulators of late ductular patterning^24^. Following processing to define the number of Seurat clusters and regress out the effects of cell cycle (Supplementary Figure 1A-C), cells clustered into five clusters using Seurat (Figure 1B). Clusters 0, 1 and 3 principally comprise of foetal hepatoblasts that continue to express a number of hepatoblast genes including *Lgr5*, *Tbx3* and *Hnf4a*. Cluster 4 are hepatocytes as defined by a number of hepatocyte markers including *Cps1*, *Cyp2d10* and *Ppara* and cluster 2 is comprised of cells that express markers of BECs, *Krt7*, *Krt19* and *Spp1* (Supplementary Figure 1D). Cells in cluster 3, express elevated levels of the master biliary transcription factor *Sox9* and the planar cell polarity genes *Vangl1* and *Vangl2* (Figure 1C). Cluster 4 also shows high *Vangl2* transcript levels. Cells within this cluster are made up from the E10.5 liver bud, prior to the initiation of definitive hepatogenesis.

**Figure 1.**
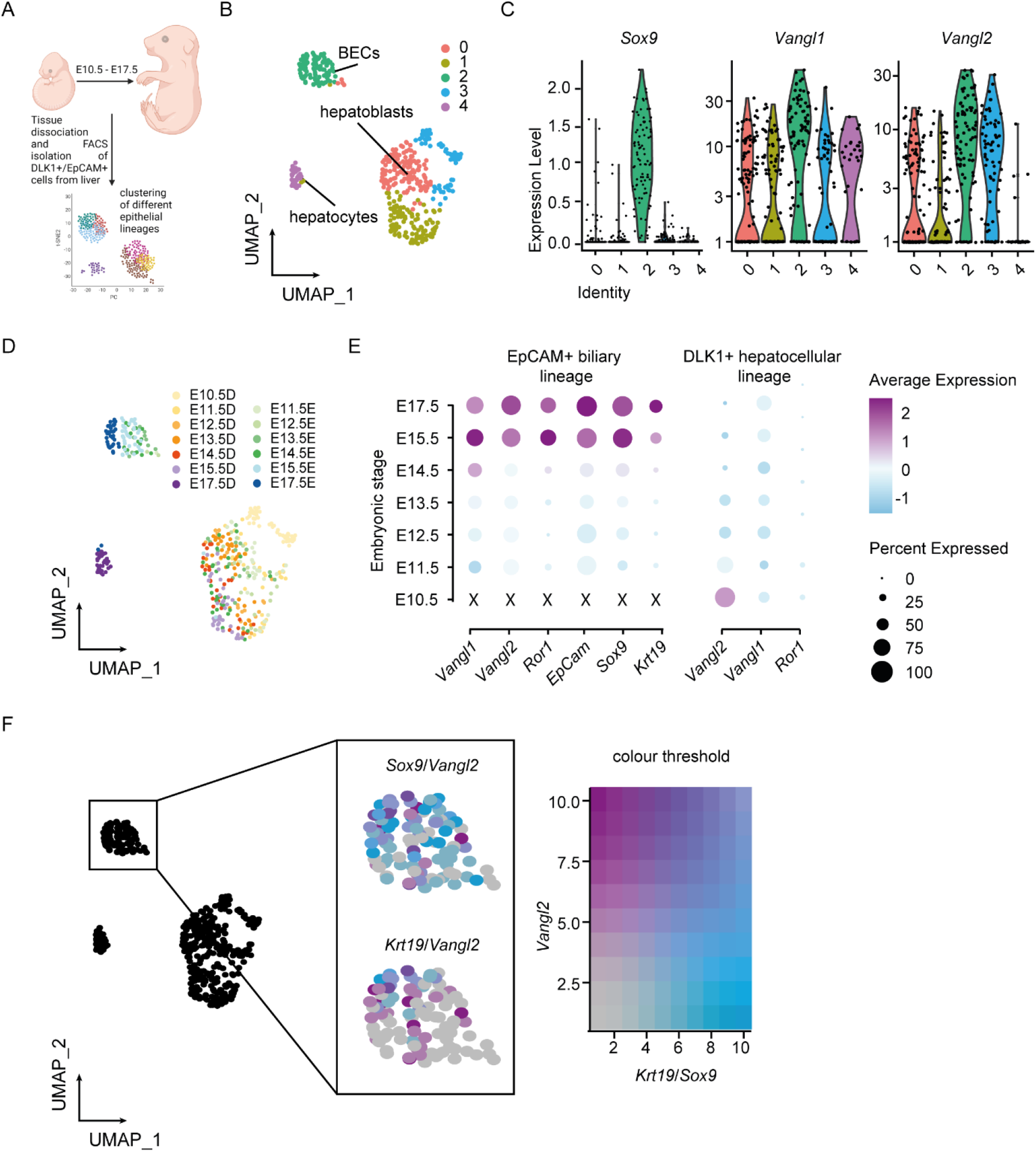
– Planar cell polarity pathway components are exclusively expressed in the biliary lineage during liver development. **A**. Schematic describing the isolation and scRNA-seq approach described by Yang et al 2017 ^24^) of E10.5-E17.5 murine liver. **B.** Seurat clustering of scRNA-seq data shows 5 distinct clusters (clusters 0-4). **C.** mRNA expression of the biliary marker, *Sox9* and core PCP components *Vangl1* and *Vangl2* between the different Seurat populations. **D.** Clustered scRNA-seq data coloured by developmental time D denotes DLK1+ cells and E EpCAM+ cells. **E.** Transcriptional expression of PCP pathway members, *Vangl1*, *Vangl2* and *Ror1* with the biliary lineage makers *EpCam*, *Sox9* and *Krt19* EPCAM+ cells (left panel) and DLK1+ cells (right panel). **F.** Correlation plot between *Sox9* and *Vangl2*, and *Krt19* and *Vangl2*.

The separation of the ductal plate and subsequent BEC (Biliary Epithelial Cell) lineage from the hepatocellular one happens at E14.5 in mice and is driven by localised signals from the portal mesenchyme^25,26^. Beyond E14.5 and following specification, ductular cells undergo further differentiation and morphogenesis. Segregating the scRNA-seq data by developmental time showed that within the BEC cluster (cluster 3) there are cells from E14.5-E17.5 (Figure 1D) indicating that this EpCAM-positive population could provide insight into the post-specification processes that govern bile duct patterning, whereas Cluster 4 (hepatocytes) was principally made up of E17.5 cells which were isolated based on DLK1. We pooled all EpCAM-positive cells or DLK1-positive cells from each developmental time point and as anticipated could see the progressive and increasing expression of biliary marker genes *Epcam*, *Sox9* and *Krt19* only within the EpCAM positive group. Similarly, *Vangl1*, *Vangl2* and *Ror1* were only transcriptionally increased within this ductular lineage and not in the DLK1-positive hepatocytes (Figure 1E).

VANGL2 is a core regulator of PCP in vertebrates and is functionally dominant over VANGL1^27^. Furthermore, ROR1 has been shown to functionally interact with VANGL2^28^. We therefore asked whether *Vangl2* expression specifically is always present in the ductular lineage or whether its expression is associated with bile duct maturation. *Vangl2* transcriptional expression does not particularly correlate with *Sox9* mRNA levels (which is expressed from the point of ductular specification onwards), however it does strongly correlate with *Krt19* expression, suggesting that *Vangl2* is intimately linked to the maturation of bile ducts as they undergo morphogenesis and is not simply present for the duration of ductulogenesis (Figure 1F).

### VANGL2 interacts with cell-cell junction proteins in BECs to pattern cell contacts

Mutations in *Vangl2* are associated with a range of ductular patterning defects across multiple organs, however, how VANGL2 results in the collective polarisation of cells and patterning of migration within a tube remains unclear. Using a transgenic mouse line which has GFP fused to the C-terminus of Vangl2 (*Vangl2^GFP^*)^29^ and whole mount FUnGI imaging we found that GFP (Green Fluorescent Protein) (and therefore VANGL2) is located at the apico-lateral membranes of Keratin-19 expressing BECs in E18.5 livers (Figure 2A). The polarisation of VANGL2 is associated with convergent extension ^19,30,31^ and as ductular morphogenesis requires the elongation of primordial ducts into a continuous biliary tree, we hypothesised that VANGL2 could coordinate the super-cellular architecture of the duct.

**Figure 2.**
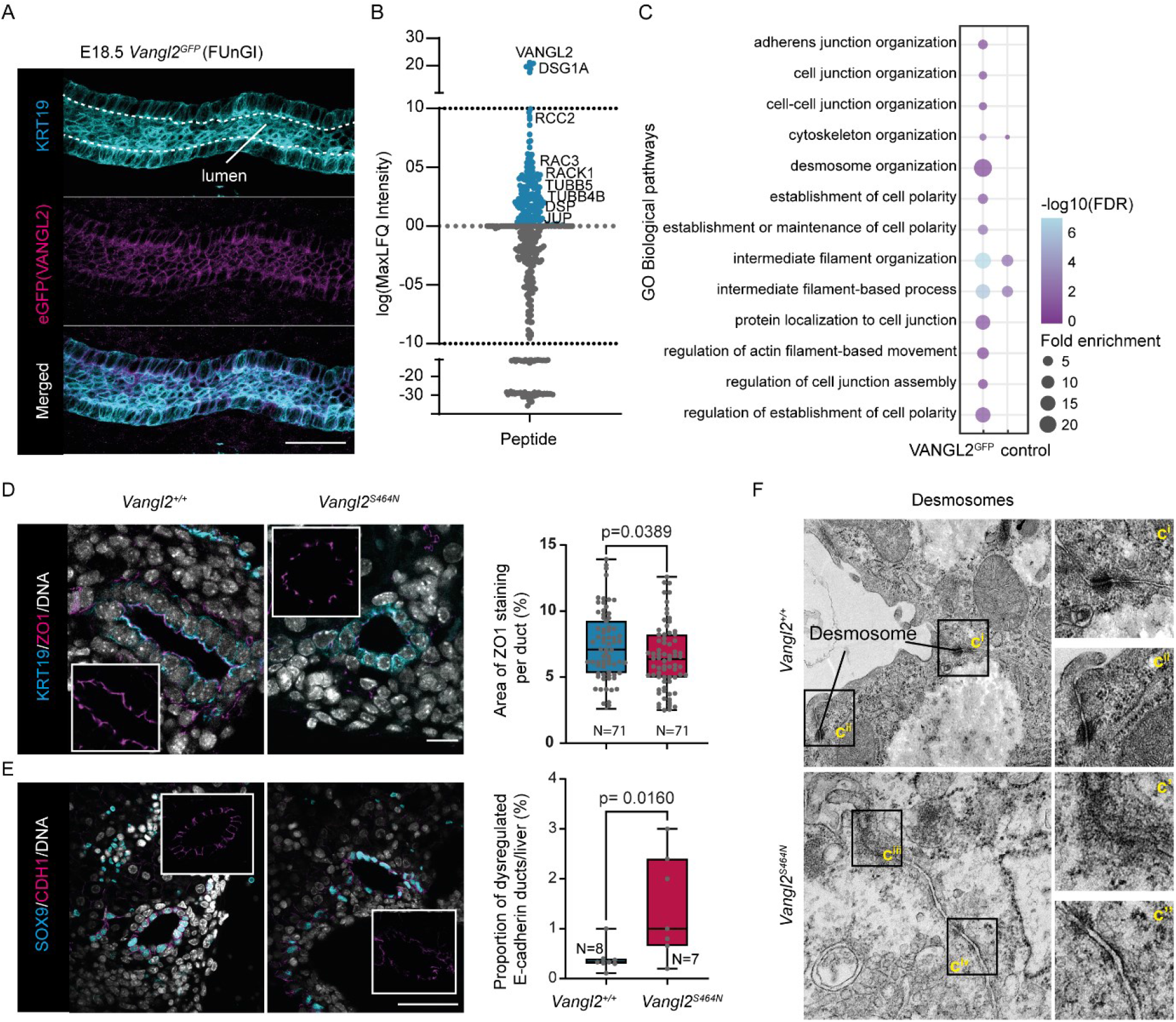
– VANGL2 interacts with cell-cell junction proteins and coordinates normal junction formation. **A**. Whole mount imaging of KRT19 positive bile duct (cyan) and GFP (magenta) in VANGL2^GFP^ bile ducts. **B.** Peptides found following co-immunoprecipitation mass spectroscopy of VANGL2^GFP^ from E18.5 livers when compared to pull down from *Vangl2^+/+^* livers (N=3 biological replicates per condition). **C.** GO Biological Pathway analysis of peptides enriched following co-immunoprecipitation mass spectroscopy **D.** Immunostaining of the biliary marker KRT19 (cyan) and tight junction protein, ZO1 (magenta) in *Vangl2^+/+^* vs *Vangl2^S464N^* livers (scale bar = 15µm). Histogram, right shows the area of ZO-1 staining within SOX9-positive cells. **E.** Immunostaining of the biliary marker SOX9 (cyan) and adherens junction protein, CDH1 (magenta) in *Vangl2^+/+^* vs *Vangl2^S464N^* livers (scale bar = 50µm), DNA in white. Histogram, shows the number of ducts with dysregulated CDH1 in *Vangl2^+/+^* vs *Vangl2^S464N^* livers. **F.** Electron micrographs of liver cells from in *Vangl2^+/+^* vs *Vangl2^S464N^* livers.

To understand this further we captured VANGL2 and its binding partners by co-immunoprecipitation of VANGL2^GFP^ from E18.5 embryonic livers and subjected these proteins to mass-spectroscopic analysis. Unsurprisingly, the top peptide we isolated was VANGL2 following GFP pulldown, however associated with this we also enriched for DSG1A, RCC2, RAC3, RACK1 and various TUBB peptides (Figure 2B and Supplementary Table 2). Furthermore, following Gene Ontology analysis of peptides that are co-precipitated with VANGL2^GFP^ we identified that amongst others, groups of peptides associated with “desmosome organisation”, “protein localisation to cell junctions” and “intermediate filament organisation” were particularly enriched (Figure 2C). It is possible that during liver development VANGL2^GFP^ is expressed by non-epithelial cell types, therefore we isolated livers from E15.5 *Vangl2^GFP^* transgenic mice and use these to derive foetal liver organoids (FLOs). FLOs are generated in a culture medium which selects for a highly purified population of biliary-lineage cells^32^. Indeed, VANGL2^GFP^ is expressed by BECs which comprise the FLOs and is physically associated with proteins involved in cell adhesion and intermediate filament organisation (Supplementary Figure 2A, 2B and Supplementary Table 2).

Collectively, these data suggested that VANGL2 can physically interact with cell junction proteins and pattern the normal formation of cell-cell contacts. Indeed, in a transgenic mouse line that carries a homozygous hypomorphic mutation in *Vangl2* (*Vangl2^S464N/S464N^*, from hereon in known as *Vangl2^S464N^*) we found significant defects in tight junctions through reduced expression and distribution of ZO-1 and Occludin (Figure 2D and Supplementary Figure 2C), adherens junctions (through deregulation of CDH1 patterning, Figure 2E) and loss of desmosomes when compared to *Vangl2^+/+^* littermate controls (Figure 2F) suggesting that loss of functional VANGL2 limits the ability of BECs to normally pattern cell-cell contacts during ductular development.

### Loss of VANGL2 functionlimits the formation of a normal biliary network

The loss of functional VANGL2 limits the normal distribution of cell-cell contacts between BECs during bile duct development (Figure 2) and, while there are no differences in overall liver size between *Vangl2^+/+^* and *Vangl2^S464N^* livers at E18.5 (Figure 3A), there is a significant reduction in the number of Keratin-19 positive ducts distributed throughout the tissue at this time point (Figure 3B). Given our data suggests that the coordination of bile duct morphogenesis by PCP proteins is a late event in liver development, we sought to determine whether the phenotypes we see at E18.5 are established earlier in ductular ontogeny or whether they are concordant with late ductular remodelling and maturation.

**Figure 3.**
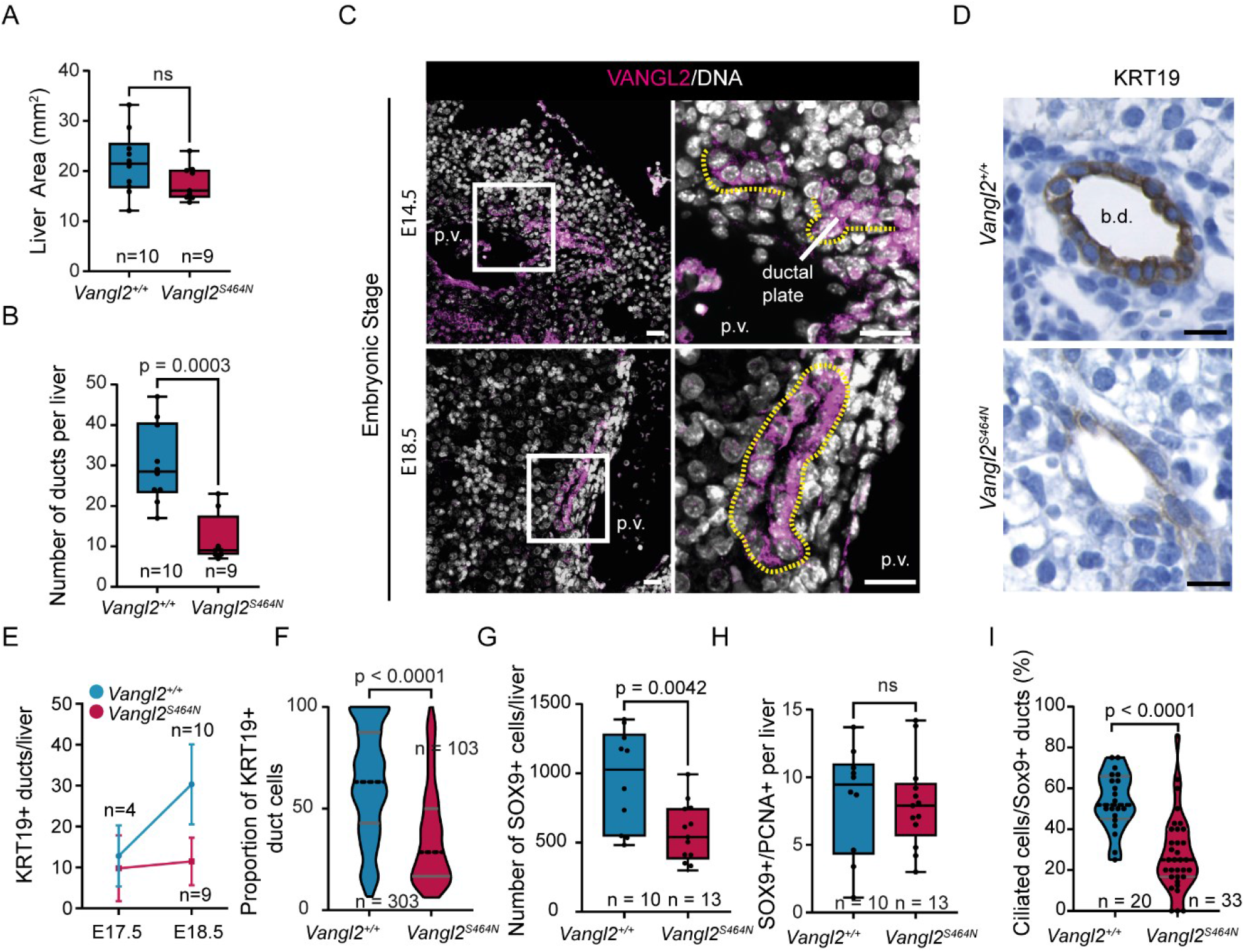
– Mice with hypomorphic *Vangl2^S464N^*do not have a normal biliary tree. **A**. Quantification of liver area in E18.5 *Vangl2^+/+^* and *Vangl2^S464N^* livers. **B.** Number of KRT19-positive bile ducts per liver in E18.5 *Vangl2^+/+^* and *Vangl2^S464N^*livers. **C.** VANGL2 immunostaining (magenta) of E14.5 ductal plate cells and E18.5 bile ducts from VANGL2^+/+^ mice (scale bar = 50µm), DNA grey. Basal surface of the cells demarcated with dotted yellow line. **D.** Immunohistochemistry for KRT19 in *Vangl2^+/+^* and *Vangl2^S464N^* livers (scale bar = 50µm). **E.** Change in the number of KRT19-positive ducts in *Vangl2^+/+^*and *Vangl2^S464N^*livers between E17.5 and E18.5. **F.** H-score (intensity) of KRT19 in ductular cells in *Vangl2^+/+^* and *Vangl2^S464N^* livers at E18.5. **G.** Total number of SOX9-positive cells and **H.** number of proliferating (PCNA-positive) SOX9-positive cells per liver. **I.** Quantification of SOX9-positive bile duct cells presenting a primary cilium (demarcated with AcTUB and ARL13B) in *Vangl2^+/+^* and *Vangl2^S464N^*.

VANGL2 is dynamically redistributed during the development of other ductular tissues, and this re-distribution is essential for the establishment of normal tissue function^8^. Upon commitment to the BEC lineage and the prior to ductular morphogenesis in the liver (at E14.5), VANGL2 is localised to the basal surface of the cells comprising the ductal plate. By E18.5, however, VANGL2 is found at the apico-lateral surface of BECs (Figure 3C), reflecting the expression pattern found with VANGL2^GFP^(Figure 2) and indicating that PCP is established by this point. Furthermore, when we quantify the differences between Keratin-19 positive bile ducts at E17.5 and E18.5 in *Vangl2^+/+^* compared to *Vangl2^S464N^* animals we found that while there are similar numbers of bile ducts between the two genotypes at E17.5 there is a substantial reduction in bile duct number by E18.5 (Figure 3D and 3E) and the number of Keratin-19 positive cells within those bile ducts is also significantly reduced (Figure 3F).

Keratin-19 is a basic, type-I Keratin that is part of the Keratin-Desmosome scaffold^33^ and which provides structural integrity to epithelial cells. It is possible then that the loss of Keratin-19 in *Vangl2^S464N^* mutant bile ducts is due to disruption of intermediate filament formation secondary to desmosome disruption. Indeed, Keratin-19 levels appear higher in *Vangl2^+/+^*livers compared to *Vangl2^S464N^*(Figure 3D). To rule this out, we immunostained *Vangl2^+/+^* or *Vangl2^S464N^* livers with SOX9 (a marker of the ductular lineage that is not associated with the cytoskeleton) and PCNA to quantify the number of proliferating biliary cells. While the number of SOX9 expressing cells was significantly reduced in *Vangl2^S464N^* mutant livers at E18.5 compared to control animals (Figure 3G) the proportion of proliferative (PCNA-positive) SOX9-positive biliary cells did not change. However, the ability of SOX9-positive cells to present a primary cilium into the lumen of the duct (as a proxy for mature BECs) was significantly impaired when *Vangl2* was mutated (Figure 3H and Supplementary Figure 2D).

### VANGL2 patterns intracellular tension and coordinates ductular connectivity

Ductular growth relies on the collective tubular migration of cells such that a primordial duct grows to the correct dimension and fuse with an adjacent duct to form a continuous structure^7,34^. To do this, cells must polarise and remodel their cytoskeletons in order that collective cell movement is coordinated. Phosphorylation of myosin light chain-2 (MLC2) results in the stabilisation of actin filaments and changes in cytoskeletal tension. In E18.5 *Vangl2^+/+^* bile ducts pMLC2^S19^ is polarised across the apical-basal axis of ductular cells with higher levels of apical pMLC2^S19^ at the apical surface. In *Vangl2^S464N^* mutant livers at the same developmental time point, however, pMLC2^S19^ is either completely absent from ductular cells or deregulated within these cells, being present at the apical, lateral and basal parts of biliary cells (Figure 4A and 4B), furthermore cells which are absent for pMLC2^S19^ are typically shorter than their wild-type counterparts (Figure 4C).

**Figure 4.**
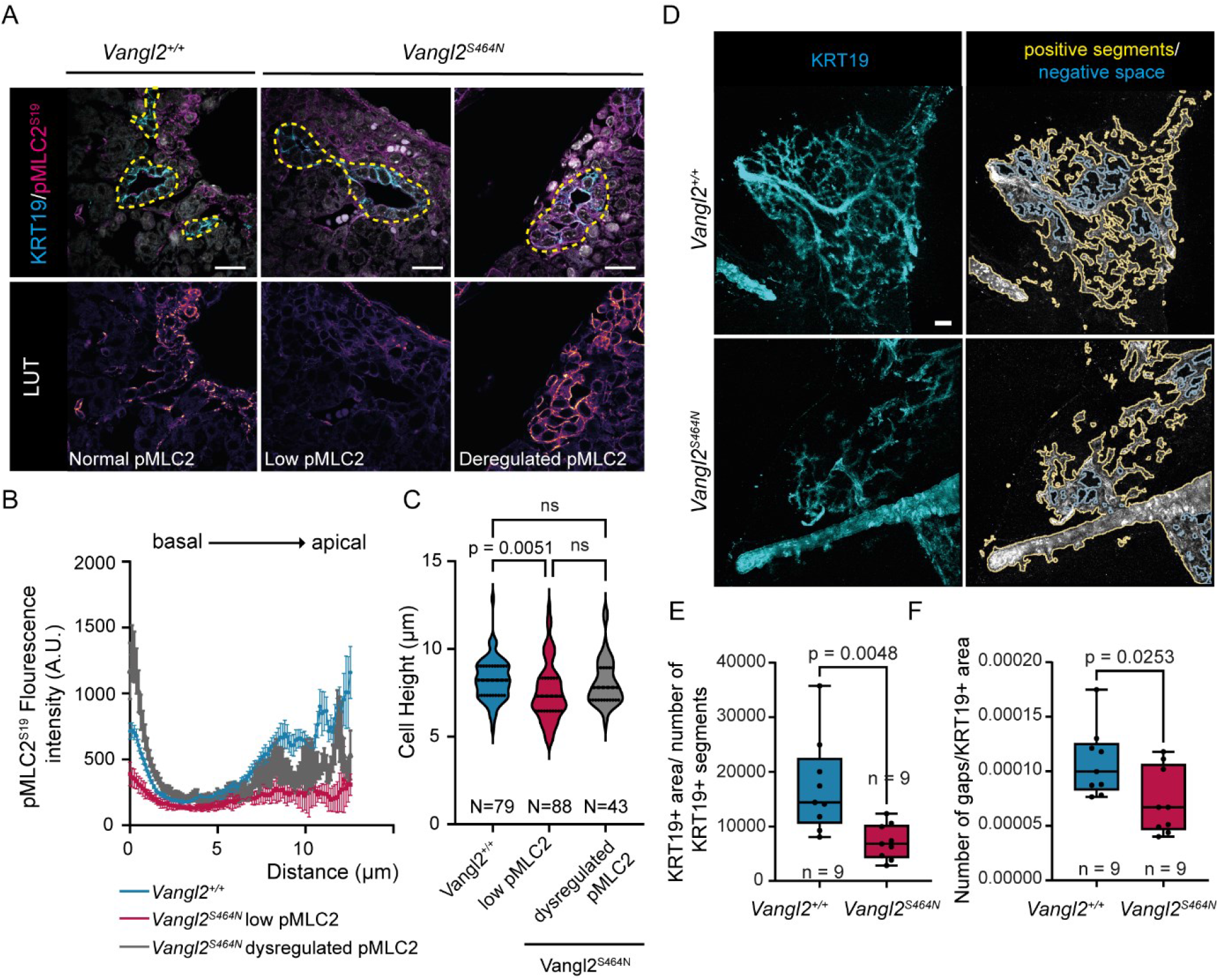
–Loss of functional VANGL2 changes the tertiary structure of bile ducts. **A**. E18.5 *Vangl2^+/+^*and *Vangl2^S464N^*livers immunostained for KRT19 (cyan) and pMLC2^S19^ (magenta), scale bar = 50µm, DNA in white. Lower panels show pMLC2^S19^ intensity. **B.** Quantification of pMLC2^S19^ signal intensity along the apico-basal axis of biliary cells in E18.5 *Vangl2^+/+^*(blue line), and in *Vangl2^S464N^*mutant animals with low pMLC2^S19^ (magenta) or dysregulated (mislocalised) pMLC2^S19^ (grey). **C.** Cell height of biliary cells from E18.5 *Vangl2^+/+^*and *Vangl2^S464N^*livers. **D.** Whole mount immunostaining for KRT19 (cyan) in *Vangl2^+/+^*and *Vangl2^S464N^*(left panels), annotations of positive segments and negative space (right panels). **E, F.** Quantification of bile duct connectivity in E18.5 *Vangl2^+/+^* and *Vangl2^S464N^*animals.

PCP-dependent patterning of the cytoskeleton is required for collective cellular migration. Using whole mount imaging of bile ducts from E18.5 livers we could demonstrate that at this stage of liver development the bile duct is formed with a complex network of small ducts connecting to a larger main duct. In *Vangl2^S464N^*embryonic livers, however, this ductular network does not form correctly, rather imaging showed that a rudimental biliary tree develops with small ductules that do not connect to each other nor do they connect to larger ducts. To quantify these phenotypic differences, we calculated the size of Keratin-19 positive segments which were significantly smaller in *Vangl2^S464N^* mice than *Vangl2^+/+^*controls (Figure 4E). In addition to smaller size ducts, we quantified the number of ducts relative to the number of gaps made by interconnecting ducts to measure “connectedness” of the biliary tree. We found that there is a significant deficiency in the connections formed between ducts, with more gaps in the ducts of *Vangl2^S464N^* mutant livers (Figure 4F).

### VANGL2 regulates planar cell polarity signalling to promote ductular morphogenesis

VANGL2 directly interacts with cell-cell junction proteins to pattern normal duct connectivity in the developing mammalian bile duct through regulation of the BEC cytoskeleton; however, whether this directly promotes the fusion of discontinuous primordial ductules to form a continuous biliary structure is difficult to assay *in vivo*. To overcome this, we isolated E15.5 livers from *Vangl2^+/+^* or *Vangl2^S464N^* embryos and following dissociation derived foetal liver organoids (FLOs) from these livers (Supplementary Figure 3A). Both *Vangl2^+/+^* and *Vangl2^S464N^* expressed equivalent levels of SOX9 and KRT19 protein (Supplementary Figure 3B) Furthermore, we found that while VANGL2 protein levels in FLOs harbouring the *Vangl2^S464N^*mutation are significantly reduced, (Figure 5A, and Supplementary Figure 3C) there is no compensation from VANGL1 (Supplementary Figure 3C). When either *Vangl2^+/+^*or *Vangl2^S464N^* FLOs are plated as single cells the organoids that form from *Vangl2^S464N^* mutant cells are significantly smaller than those from wild-type animals (Figure 5B).

**Figure 5.**
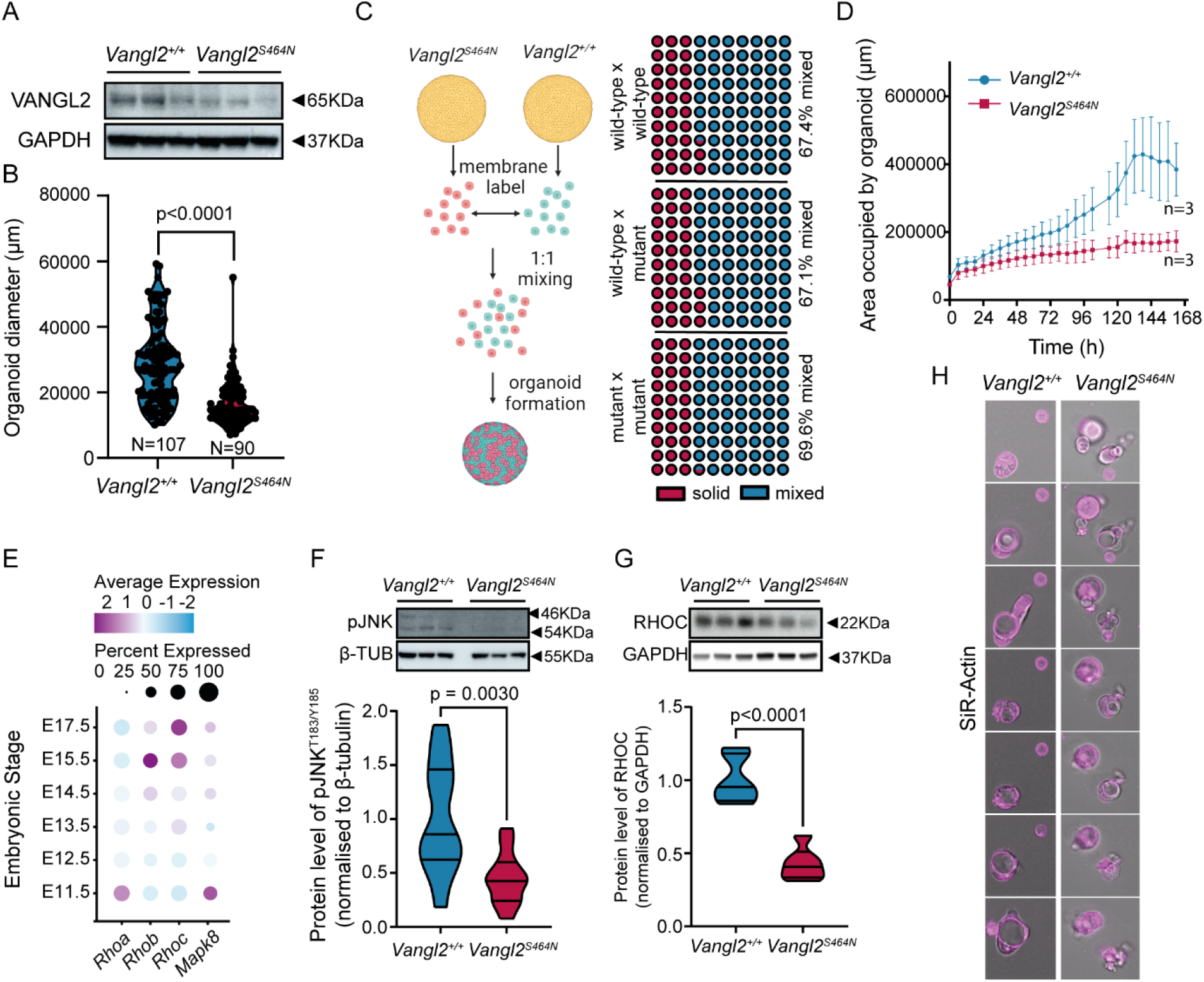
– VANGL2 promotes ductular-connectivity through actin-regulation. **A**. Immunoblot for VANGL2 and the housekeeping protein GAPDH in *Vangl2^+/+^* and *Vangl2^S464N^* organoids derived from E14.5 livers. **B.** Diameter of organoids derived from E14.5 *Vangl2^+/+^*and *Vangl2^S464N^*livers dissociated into single cells and allowed to form. **C.** Schematic and quantification of organoid admixing from *Vangl2^+/+^* and *Vangl2^S464N^* cells. Red circles denote organoids comprising of a single colour and blue circles organoids comprising two colours. (N of organoids analysed: wild-type x wildtype: 406, wild-type x Vangl2^S464N^: 420, Vangl2^S464N^ x Vangl2^S464N^:485). **D.** Growth of *Vangl2^+/+^*(blue line) *Vangl2^S464N^*(magenta line) single cells into organoids over 162h **E.** scRNA-seq from EpCAM-positive cells showing transcriptional levels of *Rhoa*, *Rhob*, *Rhoc* and *Mapk8*. **F.** Immunoblot and quantification of pJNK^T183/Y185^ in organoids derived from *Vangl2^+/+^* and *Vangl2^S464N^* E14.5 organoids. **G.** Immunoblot and quantification of RHOC in organoids derived from *Vangl2^+/+^* and *Vangl2^S464N^* E14.5 organoids. **H.** Live imaging of SiR-Actin (magenta) in E14.5 organoids derived *Vangl2^+/+^* and *Vangl2^S464N^* livers (over 24 hours).

Based on our *in vivo* data and the growth deficits seen in *Vangl2^S464N^* FLOs, we sought to determine how FLOs grow. Using time-lapse imaging over the first 6 days of organoid growth, we found that wildtype FLOs grow by forming small organoids which then fuse to form larger structures (Supplementary Movie1). We hypothesised then that small *Vangl2*-mutant organoids either fail to come together and fuse to form larger organoids or the rate of organoid fusion is significantly reduced in the *Vangl2^S464N^*-mutant. To dissect this, we dissociated either mutant or wild-type FLOs to single cells and stained these with either PKH26 or PKH67 general membrane markers. *Vangl2^+/+^*and *Vangl2^S464N^*cells were then either admixed together or admixed with themselves and the number of single colour or dual colour organoids, which was quantified (Figure 5C) to determine whether *Vangl2^S464N^*cells have an intrinsic inability to contribute to organoid formation. When FLO cells were mixed in the following combinations *Vangl2^+/+^*:*Vangl2^+/+^*, *Vangl2^+/+^*:*Vangl2^S464N^* and *Vangl2^S464N^*:*Vangl2^S464N^* we found no statistically significant differences in the ability of mutant cells to contribute to the formation of FLOs. When *Vangl2* mutant and wild-type FLO cells were plated as single cells and imaged over time, however, we found that there was a significant lag in growth of FLO derived from *Vangl2^S464N^* cells suggesting that the rate at which small organoids merge and fuse to form more substantial FLOs is limited when *Vangl2* is mutated (Figure 5D).

VANGL2 coordinates a signalling cascade which results in the activation of signalling through both ROCK/RHO^16,35^ or JNK, which itself regulates actin fibre maturation^36^. Using single cell RNAseq data from Yang et al (Figure 1) we looked at expression of the three mammalian *Rho* homologs (*Rhoa*, *Rhob* and *Rhoc*) and *Mapk8 (*the gene encoding JNK) within the EpCAM+ BEC lineage. *Rhoa* is expressed early in ductular development, however expression is lost by the time ducts are undergoing morphogenesis. *Rhob* and *Rhoc* are both expressed at the transcript level within this lineage, with increasing numbers of cells expressing *Rhoc* from E15.5. Similarly, the level of *Mapk8* is increased after ductular lineage commitment and during ductular morphogenesis (Figure 5E). In the adult regenerating bile duct JNK signalling is lost following functional *Vangl2*-loss ^37^ similarly, in FLOs the levels of pJNK^T183/Y185^ are significantly decreased (Figure 5F). Furthermore, when we specifically look for levels of RHOC we found that this is significantly reduced in FLOs derived from *Vangl2^S464N^* mutant mice (Figure 5G). Given both JNK and RHOC have a role in actin stabilisation and organisation we stained both *Vangl2^+/+^* and *Vangl2^S464N^* FLOs with the live actin stain, SiR-Actin and imaged them for 24 hours. In *Vangl2^+/+^* organoids, SiR-Actin polarises to the apical (luminal) side of the cells within the organoids as they grow and merge. In *Vangl2^S464N^*organoids, however, actin is poorly polarised, often filling the cells (Figure 5H).

The failure to connect primordial ductules together ultimately limits a duct from forming, however, its impact on normal function has not been addressed. The formation of apico-basal polarity is essential for normal ion and small molecule transport functions in BECs. We therefore treated *Vangl2^+/+^* or *Vangl2^S464N^*FLOs with Rhodamine123, a fluorescent substrate of the MDR1 transporter. In *Vangl2^+/+^* FLOs, Rhodamine123 was actively transported into the lumen of organoids and could be inhibited by co-treatment with an MDR1-inhibitor, verapamil. This was not the case in *Vangl2^S464N^* mutant FLOs, which showed a significant reduction in their ability to transport Rhodamine123 into the organoid lumen (Supplementary Figure 4A-D).

Collectively our data shows that when the function of the PCP protein VANGL2 is lost, embryonic biliary cells are no longer able to form normal cell-cell contacts and intracellular cytoplasmic tension. Failure to develop this biomechanical framework limits the rate at which primordial ducts can connect to form a complex, functional biliary network.

## Discussion

The mammalian biliary tree necessarily undergoes a number of morphological rearrangements to transition from a relatively simple epithelial sheet (which constitutes the ductal plate) into a complex, branched and continuous tubular network that follow the portal vasculature^7^. Indeed, a number of studies in mice, fish and human have shown that instructive signals from the vascular endothelium or the mesenchyme surrounding the vasculature are essential for the specification of the bile duct lineage^2,38–40^. What the post-specification signals are that regulate the formation of the biliary tree, however, have remained elusive and what mechanisms promote discontinuous, primordial ductules to elongate and intercalate to form a continuous ductular network in mammals is not clear^41^. In zebrafish, Ephrin signalling contributes to normal ductular growth and patterning^42^. Furthermore, studies using morpholinos against several components of the PCP pathway showed that these proteins are required for the formation of a normally patterned bile duct network^15^ but whether this is true in mammals and how PCP regulates bile duct development is not known.

The formation of a bile duct of the correct length and width is essential for tissue function^43^ and within the liver and other “tubular” tissues, such as the pancreas and kidney, abnormal patterning of tubules and ducts leads to organ insufficiency^44,45^. Given the essential nature of tubule and duct formation and organ function, it is unsurprising perhaps that a core group of highly conserved signals regulate this process in mammals. In tissues where tubular structures undergo classical branching morphogenesis in a highly stereotyped manner, such as the pancreas and lung ^46,47^ changes in, VANGL2 affect the ability of cells to contribute to normal tissue architecture^48^. Here, we similarly demonstrate that in the bile duct (which does not undergo classical branching morphogenesis) PCP components are transcriptionally expressed and their protein products dynamically localise to the apico-lateral membranes of BECs during ductular morphogenesis. In lung morphogenesis, *Vangl2*-abrogation results in changes in cytoskeletal mechanics^49^, however, how cellular-level changes in PCP affects super cellular patterning of tissues is less clear. We show that in addition to the classical role of VANGL2 in regulating Rho and Rac signalling (which impinges on remodelling of the cytoskeleton), VANGL2 also physically interacts with proteins that are part of the desmosome and loss of VANGL2 function results in loss or mis-localisation of cell-cell contacts, which are themselves essential for providing a group of cells collective directionality^50^.

The formation of sophisticated structures is a hallmark of tissue development. This requires the integration of chemical signals with mechanical tissue-level changes. We demonstrate for the first time that the mammalian biliary tree relies on planar cell polarity to form correctly following lineage specification and suggest that this is achieved through patterning of super-cellular tension within the duct.

## Acknowledgements

We would like to thank Matthew Pearson at the IGC (Institute of Genetics and Cancer) Advanced Imaging Resource, Lizzie Freyer at the IGC Cytometry and Single Cell Core facility. TEM was provided courtesy of the Wellcome Trust Multiuser Equipment Grant (WT104915MA) with support from Stephen Mitchell. *Funding:* MR, EC and NY are funded by an MRC Unit Award. SHW is funded by a Chief Scientist Office (CSO) Early post-doctoral fellowship (EPD/22/12). A Cancer Research UK Fellowship (C52499/A27948) funds LB. *Author Contributions:* MR planned and performed experiments, analysed data and edited the manuscript. EC, RK and NY analysed data and generated figures for the manuscript. SHW provided intellectual input, experimental design and funding for the project. LB led the project, funded the project, designed experiments, analysed data, and wrote and edited the manuscript*. Conflict of Interest:* All authors declare that they have no competing interests. *Data and materials availability:* All data is available in the manuscript or the supplementary materials. Single cell RNAseq data from this study is available from: GSE90047. All materials generated as part of this study will be made available upon request to the corresponding authors.

---

**Supplementary Figure 1:**
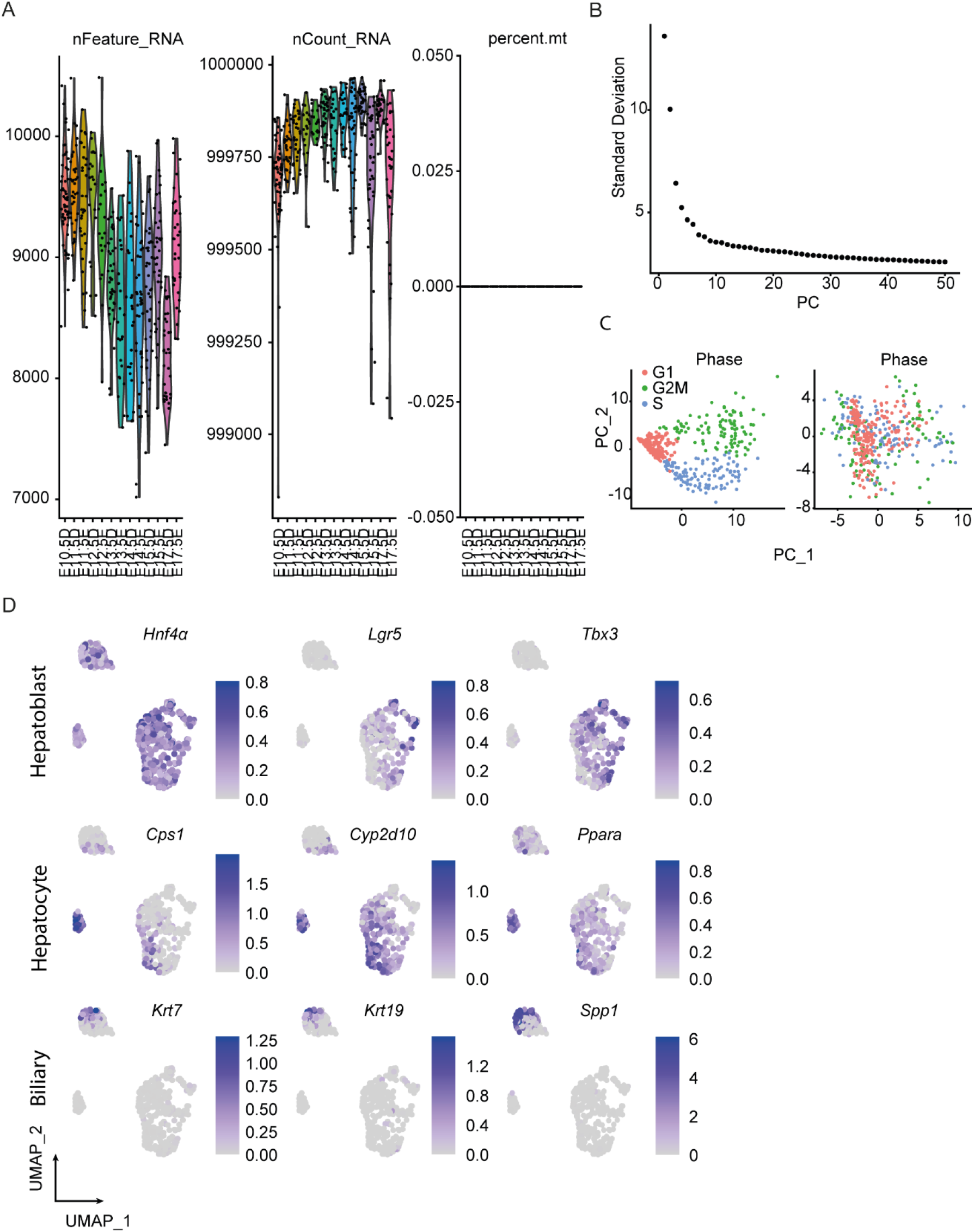
Single cell RNA sequening identifies changes in gene expression over developmental time. **A.** Single cell RNA sequencing data form Yang et al. Processed to show nFeature, nCount and mitochondrial contamination. **B.** Elbow plot to define the number of clusters used by Seurat to partition the data. **C.** Representation of the data prior to and following the regressing out of cell cycle effects. **D.** Lineage specific markers of Hepatoblasts (*Hnf4α*, *Lgr5* and *Tbx3*), Hepatocytes (*Cps1*, *Cyp2d10* and *Ppara*) and BECs (*Krt7*, *Krt19* and *Spp1*).

**Supplementary Figure 2:**
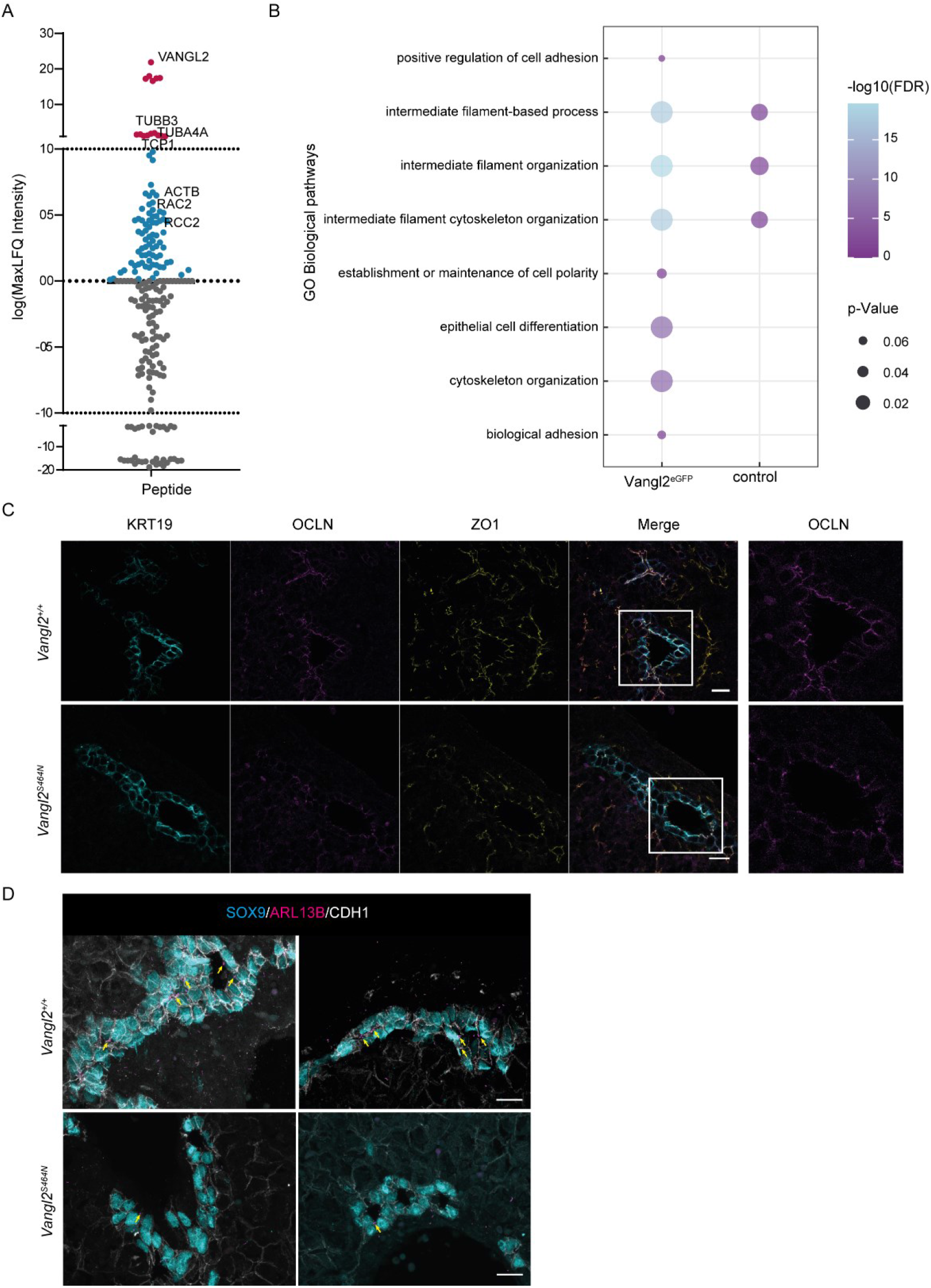
VANGL2 is asociated with cell-cell juntional stability in BECs. **A.** Significantly enriched peptides following VANGL2^GFP^ pulldown and mass spectroscopy from *Vangl2^GFP^* FLOs compared to *Vangl2^+/+^* FLOs. **B.** Enriched GOTerms for peptides isolated from co-IP mass spectrometry of VANGL2^GFP^ bait. **C.** Immunoflourescent staining of BECs with KRT19 (cyan) and TJ proteins OCLN (magenta) and ZO1 (yellow) in *Vangl2^+/+^*and *Vangl2^S464N^* livers at E18.5. Scale bar = 20µm. Insets identify regions which are shown in right panels. **D.** Immunoflourescent staining of E18.5 SOX9-positive biliary cells (cyan) from *Vangl2^+/+^*and *Vangl2^S464N^*stained for the AJ protein, CDH1 (white) and the primary cilia marker, ARL13B (magenta). Scale bar = 20µm.

**Supplementary Figure 3:**
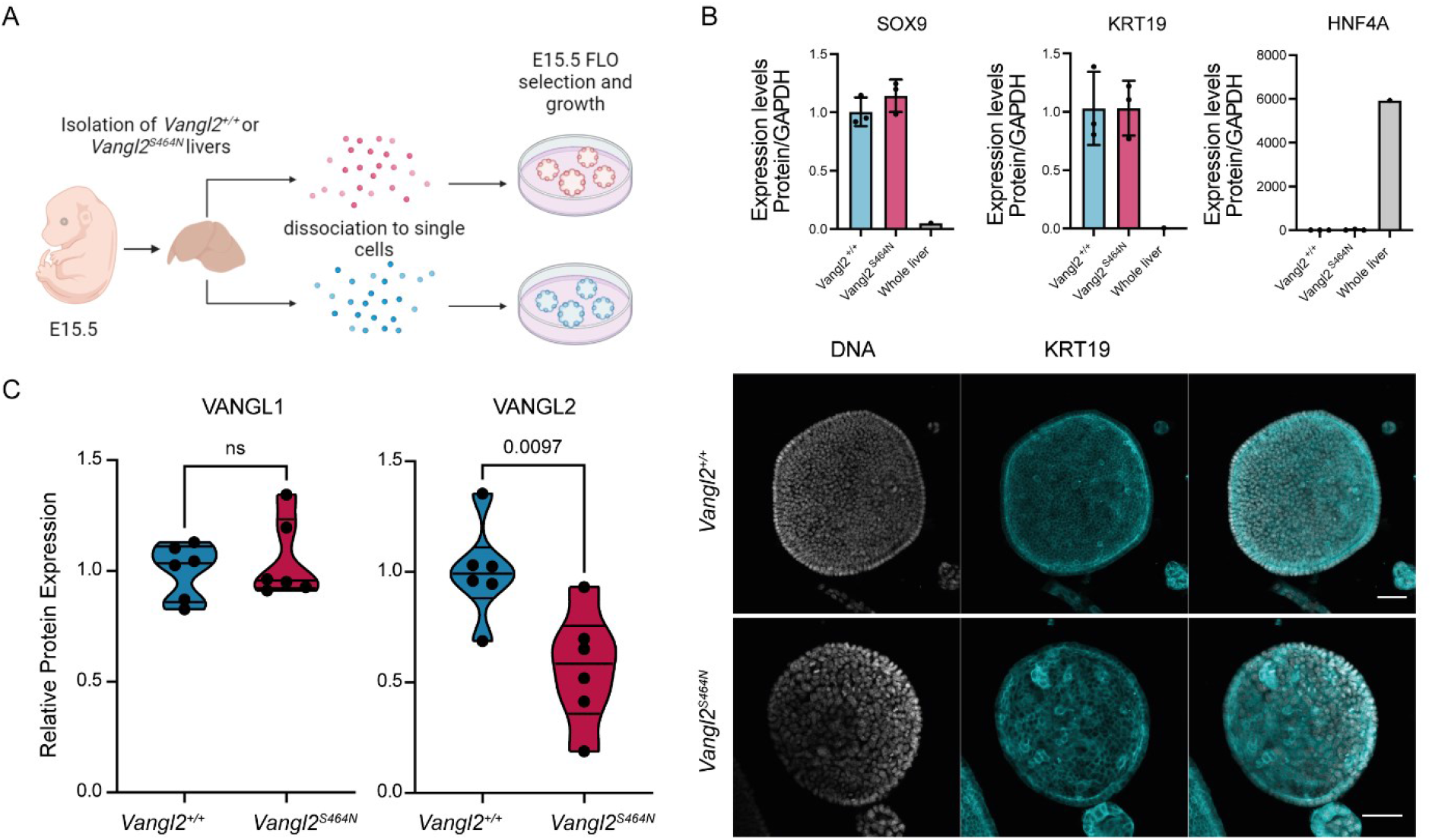
Generating FLOs from Vangl2^S464N^ mutant livers. **A.** A schematic representation of the derivation of *Vangl2^+/+^*and *Vangl2^S464N^*FLOs. **B.** mRNA expression of *Sox9*, *Krt19* and *Hnf4a* (upper panels) in *Vangl2^+/+^* and *Vangl2^S464N^* FLOs compared to whole wild-type liver (N=3 per group, except whole liver where N=1). Lower panels show immunostaining for KRT19 (cyan) in *Vangl2^+/+^* and *Vangl2^S464N^* FLOs. DNA is represented in white. Scale bar = 50µm. **C.** Quantification of immunoblots for VANGL1 and VANGL2 from proteins isolated from *Vangl2^+/+^* and *Vangl2^S464N^* FLOs (N=6).

**Supplementary Figure 4:**
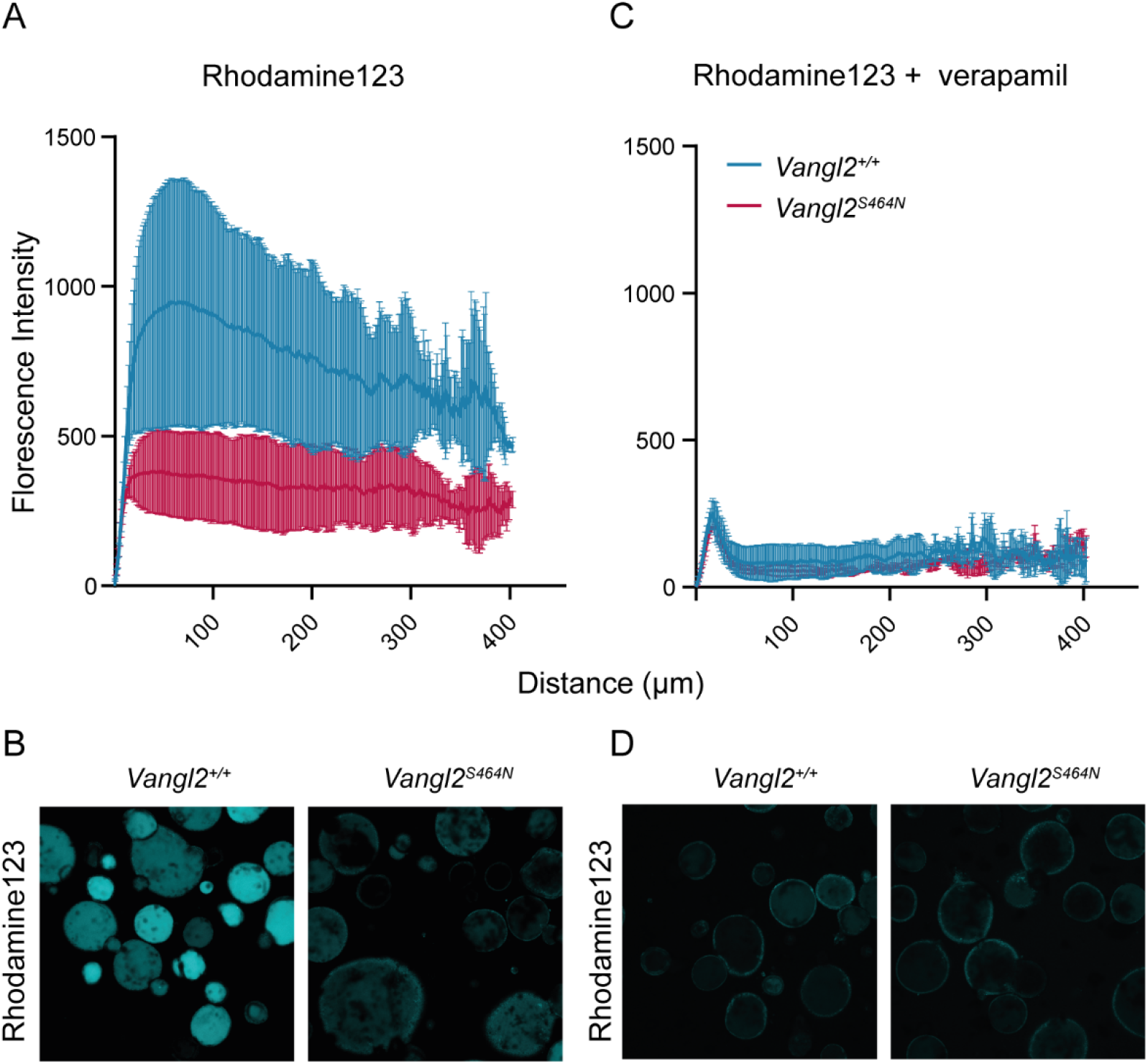
Loss of VANGL2 limits apico-basal transport in FLOs. **A.** Quantification of Rhodamine123 intensity in *Vangl2^+/+^* and *Vangl2^S464N^* organoids shown in **B. C.** Quantification of Rhodamine123 intensity in *Vangl2^+/+^* and *Vangl2^S464N^* organoids shown in **D.** Following treatment with the MDR1 inhibitor, Verapamil.

