## Supplementary Table 1 for "Planar cell polarity is essential for the architectural patterning of the mammalian biliary tree"

| **Protein ID** | **Gene** | **Organism** | **Description** | **V2GFP_log** | **WT_log** | **Subtraction value** |
| --- | --- | --- | --- | --- | --- | --- |
| Q91ZD4 | Vangl2 | Mus musculus | Vang-like protein 2 | 21.1094883 | 0 | 21.10948827 |
| P01868 | Ighg1 | Mus musculus | Ig gamma-1 chain C region secreted form | 20.7860304 | 0 | 20.78603043 |
| A0A075B5R5 | Ighv4-1 | Mus musculus | Immunoglobulin heavy variable 4-1 (Fragment) | 20.6082333 | 0 | 20.60823325 |
| P01675 |  | Mus musculus | Ig kappa chain V-VI region XRPC 44 | 20.5668259 | 0 | 20.56682593 |
| Q7TNV0 | Dek | Mus musculus | Protein DEK | 20.0115445 | 0 | 20.01154449 |
| Q61495 | Dsg1a | Mus musculus | Desmoglein-1-alpha | 19.6154679 | 0 | 19.61546789 |
| P62911 | Rpl32 | Mus musculus | 60S ribosomal protein L32 | 19.5032949 | 0 | 19.50329488 |
| P26350 | Ptma | Mus musculus | Prothymosin alpha | 18.8200058 | 0 | 18.82000577 |
| Q9JKV1 | Adrm1 | Mus musculus | Proteasomal ubiquitin receptor ADRM1 | 17.5066005 | 0 | 17.50660047 |
| A0A075B5U4 | Ighv1-18 | Mus musculus | Immunoglobulin heavy variable V1-18 | 21.8956521 | 20.5764078 | 1.319244357 |
| O88569-2 | Hnrnpa2b1 | Mus musculus | Isoform 2 of Heterogeneous nuclear ribonucleoproteins A2/B1 | 19.7491669 | 18.6666294 | 1.082537586 |
| Q501J6 | Ddx17 | Mus musculus | Probable ATP-dependent RNA helicase DDX17 | 19.7042283 | 18.6260291 | 1.078199201 |
| Q61656 | Ddx5 | Mus musculus | Probable ATP-dependent RNA helicase DDX5 | 19.7042283 | 18.6260291 | 1.078199201 |
| E9Q6E5 | Srsf11 | Mus musculus | Serine and arginine-rich-splicing factor 11 | 21.3469708 | 20.2762504 | 1.07072044 |
| P19253 | Rpl13a | Mus musculus | 60S ribosomal protein L13a | 23.5754248 | 22.5074769 | 1.067947902 |
| Q5SUF2-2 | Luc7l3 | Mus musculus | Isoform 2 of Luc7-like protein 3 | 20.0194212 | 19.0233738 | 0.996047354 |
| Q8BK67 | Rcc2 | Mus musculus | Protein RCC2 | 19.6957727 | 18.7285703 | 0.967202453 |
| Q99JB2 | Stoml2 | Mus musculus | Stomatin-like protein 2, mitochondrial | 19.5129836 | 18.5833644 | 0.929619199 |
| P67778 | Phb | Mus musculus | Prohibitin | 22.8349995 | 21.9243276 | 0.910671901 |
| A0A0B4J1H7 | Igkv1-135 | Mus musculus | Immunoglobulin kappa variable 1-135 (Fragment) | 20.8912783 | 20.0331749 | 0.858103398 |
| O35129 | Phb2 | Mus musculus | Prohibitin-2 | 23.1749892 | 22.3966633 | 0.7783259 |
| P40142 | Tkt | Mus musculus | Transketolase | 20.3403628 | 19.6162669 | 0.724095873 |
| Q62376 | Snrnp70 | Mus musculus | U1 small nuclear ribonucleoprotein 70 kDa | 20.1123117 | 19.3943788 | 0.717932893 |
| P02104 | Hbb-y | Mus musculus | Hemoglobin subunit epsilon-Y2 | 25.1794961 | 24.5105073 | 0.6689888 |
| Q9D823 | Rpl37 | Mus musculus | 60S ribosomal protein L37 | 20.5019363 | 19.8892529 | 0.612683377 |
| Q9D8E6 | Rpl4 | Mus musculus | 60S ribosomal protein L4 | 21.953032 | 21.3451994 | 0.607832653 |
| A0A0J9YUZ4 | Hmgb1 | Mus musculus | High mobility group protein 1 (Fragment) | 19.8013002 | 19.193508 | 0.607792203 |
| P69905 | HBA1 | Homo sapiens | Hemoglobin subunit alpha | 23.7593876 | 23.1725493 | 0.586838309 |
| P60764 | Rac3 | Mus musculus | Ras-related C3 botulinum toxin substrate 3 | 20.746132 | 20.1739705 | 0.572161482 |
| P06467 | Hbz | Mus musculus | Hemoglobin subunit zeta | 23.7389235 | 23.1707446 | 0.568178862 |
| A8DUK4 | Hbb-bs | Mus musculus | Beta-globin | 25.1718829 | 24.6208677 | 0.551015169 |
| P80318 | Cct3 | Mus musculus | T-complex protein 1 subunit gamma | 19.740877 | 19.1997931 | 0.541083878 |
| P61255 | Rpl26 | Mus musculus | 60S ribosomal protein L26 | 20.9418769 | 20.4036791 | 0.538197801 |
| Q922Q8 | Lrrc59 | Mus musculus | Leucine-rich repeat-containing protein 59 | 19.6622316 | 19.1467815 | 0.515450112 |
| Q8BX70-3 | Vps13c | Mus musculus | Isoform 3 of Vacuolar protein sorting-associated protein 13C | 23.1346311 | 22.6322303 | 0.502400754 |
| P14115 | Rpl27a | Mus musculus | 60S ribosomal protein L27a | 22.622546 | 22.1215416 | 0.501004403 |
| P84104-2 | Srsf3 | Mus musculus | Isoform Short of Serine/arginine-rich splicing factor 3 | 22.9224568 | 22.4252605 | 0.497196354 |
| P47963 | Rpl13 | Mus musculus | 60S ribosomal protein L13 | 22.0191071 | 21.5225565 | 0.496550574 |
| P14148 | Rpl7 | Mus musculus | 60S ribosomal protein L7 | 22.1419009 | 21.6554565 | 0.486444376 |
| P02301 | H3-5 | Mus musculus | Histone H3.3C | 24.4234217 | 23.9405574 | 0.482864313 |
| Q6NXH9 | Krt73 | Mus musculus | Keratin, type II cytoskeletal 73 | 27.7453498 | 27.2802967 | 0.465053037 |
| P62267 | Rps23 | Mus musculus | 40S ribosomal protein S23 | 22.9897098 | 22.5270722 | 0.462637553 |
| Q91VB8 | Hba-a1 | Mus musculus | Alpha globin 1 | 25.8117643 | 25.3511075 | 0.460656837 |
| P84099 | Rpl19 | Mus musculus | 60S ribosomal protein L19 | 24.5105073 | 24.0690721 | 0.441435189 |
| Q9D3D9 | Atp5f1d | Mus musculus | ATP synthase subunit delta, mitochondrial | 20.8336831 | 20.3982661 | 0.435417088 |
| P62827 | Ran | Mus musculus | GTP-binding nuclear protein Ran | 22.9734816 | 22.5396686 | 0.433812981 |
| Q9CXW4 | Rpl11 | Mus musculus | 60S ribosomal protein L11 | 22.4919398 | 22.0581695 | 0.433770329 |
| P49717 | Mcm4 | Mus musculus | DNA replication licensing factor MCM4 | 18.5552174 | 18.1216662 | 0.433551237 |
| Q60634 | Flot2 | Mus musculus | Flotillin-2 | 22.6650069 | 22.2331091 | 0.431897765 |
| A0A3B2WBL1 | Rpl10a | Mus musculus | Ribosomal protein | 21.0214641 | 20.5973035 | 0.424160577 |
| P60843 | Eif4a1 | Mus musculus | Eukaryotic initiation factor 4A-I | 22.4131756 | 21.9914123 | 0.42176324 |
| P11404 | Fabp3 | Mus musculus | Fatty acid-binding protein, heart | 21.2917744 | 20.8729592 | 0.418815196 |
| Q9D883 | U2af1 | Mus musculus | Splicing factor U2AF 35 kDa subunit | 21.351675 | 20.9376141 | 0.414060939 |
| Q7TNC4-4 | Luc7l2 | Mus musculus | Isoform 4 of Putative RNA-binding protein Luc7-like 2 | 22.7070346 | 22.299958 | 0.407076607 |
| P68040 | Rack1 | Mus musculus | Receptor of activated protein C kinase 1 | 20.0173288 | 19.6139883 | 0.403340442 |
| P80317 | Cct6a | Mus musculus | T-complex protein 1 subunit zeta | 19.5627801 | 19.1600304 | 0.402749642 |
| P52480-2 | Pkm | Mus musculus | Isoform M1 of Pyruvate kinase PKM | 20.7469238 | 20.3448011 | 0.40212274 |
| Q6PDM2 | Srsf1 | Mus musculus | Serine/arginine-rich splicing factor 1 | 20.5678427 | 20.1673772 | 0.400465429 |
| P04264 | KRT1 | Homo sapiens | Keratin, type II cytoskeletal 1 | 27.9088485 | 27.5089974 | 0.399851095 |
| P02089 | Hbb-b2 | Mus musculus | Hemoglobin subunit beta-2 | 24.9890188 | 24.592634 | 0.396384792 |
| Q8BP67 | Rpl24 | Mus musculus | 60S ribosomal protein L24 | 22.1833641 | 21.7901888 | 0.393175364 |
| P60867 | Rps20 | Mus musculus | 40S ribosomal protein S20 | 21.8023499 | 21.4104758 | 0.391874127 |
| Q6NV83-2 | U2surp | Mus musculus | Isoform 2 of U2 snRNP-associated SURP motif-containing protein | 22.2110616 | 21.8279927 | 0.383068836 |
| Q61781 | Krt14 | Mus musculus | Keratin, type I cytoskeletal 14 | 26.6458141 | 26.2731986 | 0.372615508 |
| O08917 | Flot1 | Mus musculus | Flotillin-1 | 23.1410159 | 22.7684096 | 0.372606257 |
| P62245 | Rps15a | Mus musculus | 40S ribosomal protein S15a | 22.5838457 | 22.2158046 | 0.368041052 |
| P62754 | Rps6 | Mus musculus | 40S ribosomal protein S6 | 23.7285815 | 23.364528 | 0.364053571 |
| Q921F2 | Tardbp | Mus musculus | TAR DNA-binding protein 43 | 20.1575068 | 19.8084685 | 0.349038267 |
| Q8VH51-2 | Rbm39 | Mus musculus | Isoform 2 of RNA-binding protein 39 | 20.765705 | 20.418013 | 0.347692057 |
| Q3UEB3-2 | Puf60 | Mus musculus | Isoform 2 of Poly(U)-binding-splicing factor PUF60 | 19.9692632 | 19.622377 | 0.346886189 |
| Q68FD5 | Cltc | Mus musculus | Clathrin heavy chain 1 | 20.210849 | 19.8668104 | 0.344038575 |
| Q99LX0 | Park7 | Mus musculus | Parkinson disease protein 7 homolog | 20.8941967 | 20.5511633 | 0.343033371 |
| P62806 | H4c1 | Mus musculus | Histone H4 | 24.323886 | 23.9846799 | 0.339206086 |
| Q99KI0 | Aco2 | Mus musculus | Aconitate hydratase, mitochondrial | 19.5712898 | 19.2371943 | 0.334095522 |
| P02088 | Hbb-b1 | Mus musculus | Hemoglobin subunit beta-1 | 24.9583686 | 24.6264488 | 0.331919867 |
| Q6ZWZ4 | Rpl36 | Mus musculus | 60S ribosomal protein L36 | 20.6751182 | 20.3444768 | 0.330641402 |
| P62852 | Rps25 | Mus musculus | 40S ribosomal protein S25 | 20.8365008 | 20.5109671 | 0.325533639 |
| Q9JKX6 | Nudt5 | Mus musculus | ADP-sugar pyrophosphatase | 19.3222952 | 18.9999972 | 0.322298072 |
| P41105 | Rpl28 | Mus musculus | 60S ribosomal protein L28 | 21.4362201 | 21.1195903 | 0.316629824 |
| Q9D7S7-2 | Rpl22l1 | Mus musculus | Isoform 2 of 60S ribosomal protein L22-like 1 | 20.1962527 | 19.886942 | 0.309310704 |
| P62889 | Rpl30 | Mus musculus | 60S ribosomal protein L30 | 20.0048362 | 19.6983782 | 0.306457992 |
| P06151 | Ldha | Mus musculus | L-lactate dehydrogenase A chain | 21.5407085 | 21.2350829 | 0.305625575 |
| P16381 | D1Pas1 | Mus musculus | Putative ATP-dependent RNA helicase Pl10 | 20.5858928 | 20.2841188 | 0.30177408 |
| P43277 | H1-3 | Mus musculus | Histone H1.3 | 21.5964415 | 21.3099598 | 0.286481698 |
| Q9D1R9 | Rpl34 | Mus musculus | 60S ribosomal protein L34 | 20.69112 | 20.4046776 | 0.286442455 |
| P42932 | Cct8 | Mus musculus | T-complex protein 1 subunit theta | 19.8073707 | 19.5289178 | 0.278452927 |
| Q6IFX2 | Krt42 | Mus musculus | Keratin, type I cytoskeletal 42 | 26.6458141 | 26.3711917 | 0.27462238 |
| P57776-2 | Eef1d | Mus musculus | Isoform 2 of Elongation factor 1-delta | 19.8148911 | 19.5415912 | 0.273299873 |
| P47915 | Rpl29 | Mus musculus | 60S ribosomal protein L29 | 21.774555 | 21.50304 | 0.271514996 |
| P35980 | Rpl18 | Mus musculus | 60S ribosomal protein L18 | 21.8123615 | 21.5427129 | 0.269648663 |
| Q61753 | Phgdh | Mus musculus | D-3-phosphoglycerate dehydrogenase | 20.6758894 | 20.4105105 | 0.26537897 |
| P68372 | Tubb4b | Mus musculus | Tubulin beta-4B chain | 23.4676215 | 23.2023564 | 0.265265043 |
| P35527 | KRT9 | Homo sapiens | Keratin, type I cytoskeletal 9 | 26.5897801 | 26.3256025 | 0.264177587 |
| P99024 | Tubb5 | Mus musculus | Tubulin beta-5 chain | 23.4676215 | 23.2049508 | 0.262670694 |
| Q8VEM8 | Slc25a3 | Mus musculus | Phosphate carrier protein, mitochondrial | 20.5272537 | 20.2659819 | 0.261271798 |
| P63260 | Actg1 | Mus musculus | Actin, cytoplasmic 2 | 25.0105199 | 24.7593876 | 0.251132317 |
| Q64523 | H2ac20 | Mus musculus | Histone H2A type 2-C | 24.2316923 | 23.9846799 | 0.247012388 |
| P48678 | Lmna | Mus musculus | Prelamin-A/C | 20.7286903 | 20.4824987 | 0.246191638 |
| Q91Z53 | Grhpr | Mus musculus | Glyoxylate reductase/hydroxypyruvate reductase | 20.7550005 | 20.5114243 | 0.243576174 |
| Q8VEK3-2 | Hnrnpu | Mus musculus | Isoform 2 of Heterogeneous nuclear ribonucleoprotein U | 19.7375302 | 19.496257 | 0.241273296 |
| P61750 | Arf4 | Mus musculus | ADP-ribosylation factor 4 | 20.5792174 | 20.3402272 | 0.238990227 |
| Q3TWW8 | Srsf6 | Mus musculus | Serine/arginine-rich splicing factor 6 | 19.7816027 | 19.5431804 | 0.238422281 |
| P60710 | Actb | Mus musculus | Actin, cytoplasmic 1 | 25.0105199 | 24.7745474 | 0.23597251 |
| P10854 | H2bc14 | Mus musculus | Histone H2B type 1-M | 25.3169996 | 25.0853739 | 0.231625701 |
| Q61881 | Mcm7 | Mus musculus | DNA replication licensing factor MCM7 | 19.6188209 | 19.3875968 | 0.231224111 |
| P43274 | H1-4 | Mus musculus | Histone H1.4 | 21.4175322 | 21.1871266 | 0.230405599 |
| P0CG49 | Ubb | Mus musculus | Polyubiquitin-B | 20.9046905 | 20.676406 | 0.22828457 |
| Q8BL97-2 | Srsf7 | Mus musculus | Isoform 2 of Serine/arginine-rich splicing factor 7 | 20.6408754 | 20.4129692 | 0.227906242 |
| Q9CPR4 | Rpl17 | Mus musculus | 60S ribosomal protein L17 | 20.6953727 | 20.4720105 | 0.223362127 |
| P08003 | Pdia4 | Mus musculus | Protein disulfide-isomerase A4 | 19.8610765 | 19.6470802 | 0.213996208 |
| P07901 | Hsp90aa1 | Mus musculus | Heat shock protein HSP 90-alpha | 21.9651057 | 21.752533 | 0.212572706 |
| Q9DCW4 | Etfb | Mus musculus | Electron transfer flavoprotein subunit beta | 19.9369764 | 19.7252233 | 0.211753094 |
| Q569Z5 | Ddx46 | Mus musculus | Probable ATP-dependent RNA helicase DDX46 | 20.1736352 | 19.9653598 | 0.208275378 |
| P67984 | Rpl22 | Mus musculus | 60S ribosomal protein L22 | 22.3450828 | 22.139524 | 0.20555874 |
| P11983 | Tcp1 | Mus musculus | T-complex protein 1 subunit alpha | 19.9342779 | 19.7293643 | 0.204913524 |
| Q8BTM8 | Flna | Mus musculus | Filamin-A | 19.6633157 | 19.4591896 | 0.204126175 |
| P62900 | Rpl31 | Mus musculus | 60S ribosomal protein L31 | 22.1544394 | 21.9534946 | 0.200944812 |
| Q9Z2I9 | Sucla2 | Mus musculus | Succinate--CoA ligase [ADP-forming] subunit beta, mitochondrial | 19.8422444 | 19.6437392 | 0.198505103 |
| P62242 | Rps8 | Mus musculus | 40S ribosomal protein S8 | 22.5052639 | 22.3072244 | 0.198039422 |
| P0C0S6 | H2az1 | Mus musculus | Histone H2A.Z | 24.2606922 | 24.0690721 | 0.191620073 |
| Q60605-2 | Myl6 | Mus musculus | Isoform Smooth muscle of Myosin light polypeptide 6 | 21.81639 | 21.6315656 | 0.184824314 |
| P62737 | Acta2 | Mus musculus | Actin, aortic smooth muscle | 24.1094864 | 23.9315686 | 0.177917792 |
| Q8BHD7-2 | Ptbp3 | Mus musculus | Isoform 2 of Polypyrimidine tract-binding protein 3 | 20.254694 | 20.0785742 | 0.176119758 |
| P14869 | Rplp0 | Mus musculus | 60S acidic ribosomal protein P0 | 20.7968294 | 20.6219095 | 0.174919919 |
| C0HKE1 | H2ac4 | Mus musculus | Histone H2A type 1-B | 24.267852 | 24.0934563 | 0.174395705 |
| P63101 | Ywhaz | Mus musculus | 14-3-3 protein zeta/delta | 21.0226093 | 20.8482591 | 0.17435013 |
| Q6ZWV3 | Rpl10 | Mus musculus | 60S ribosomal protein L10 | 21.8039127 | 21.6320569 | 0.171855844 |
| Q922D8 | Mthfd1 | Mus musculus | C-1-tetrahydrofolate synthase, cytoplasmic | 20.5354093 | 20.3641383 | 0.171271039 |
| Q9R0H5 | Krt71 | Mus musculus | Keratin, type II cytoskeletal 71 | 27.3409595 | 27.1699733 | 0.170986197 |
| Q9CX86 | Hnrnpa0 | Mus musculus | Heterogeneous nuclear ribonucleoprotein A0 | 19.552869 | 19.3854148 | 0.167454175 |
| P17182 | Eno1 | Mus musculus | Alpha-enolase | 20.6999909 | 20.5325479 | 0.16744302 |
| P60335 | Pcbp1 | Mus musculus | Poly(rC)-binding protein 1 | 21.6971276 | 21.5309973 | 0.166130288 |
| P02662 | CSN1S1 | Bos taurus | Alpha-S1-casein | 24.7695118 | 24.6039939 | 0.1655179 |
| P45952 | Acadm | Mus musculus | Medium-chain specific acyl-CoA dehydrogenase, mitochondrial | 19.0765788 | 18.9111678 | 0.16541097 |
| Q9DB20 | Atp5po | Mus musculus | ATP synthase subunit O, mitochondrial | 20.5391741 | 20.3771069 | 0.162067223 |
| P10126 | Eef1a1 | Mus musculus | Elongation factor 1-alpha 1 | 24.7644586 | 24.6096405 | 0.154818109 |
| Q61990-2 | Pcbp2 | Mus musculus | Isoform 2 of Poly(rC)-binding protein 2 | 21.3475488 | 21.1927559 | 0.154792949 |
| P63017 | Hspa8 | Mus musculus | Heat shock cognate 71 kDa protein | 21.905256 | 21.7547635 | 0.150492468 |
| P47911 | Rpl6 | Mus musculus | 60S ribosomal protein L6 | 21.517826 | 21.3707821 | 0.147043919 |
| Q8VDD5 | Myh9 | Mus musculus | Myosin-9 | 21.863245 | 21.7170091 | 0.146235951 |
| E9Q557 | Dsp | Mus musculus | Desmoplakin | 20.6647131 | 20.5211178 | 0.143595271 |
| Q6ZWN5 | Rps9 | Mus musculus | 40S ribosomal protein S9 | 22.7253854 | 22.5924982 | 0.132887185 |
| Q6ZWY3 | Rps27l | Mus musculus | 40S ribosomal protein S27-like | 21.6403069 | 21.508328 | 0.131978996 |
| P29341 | Pabpc1 | Mus musculus | Polyadenylate-binding protein 1 | 19.1153868 | 18.9892312 | 0.126155554 |
| Q8K194-2 | Snrnp27 | Mus musculus | Isoform 2 of U4/U6.U5 small nuclear ribonucleoprotein 27 kDa protein | 20.2255927 | 20.0999668 | 0.125625898 |
| Q8BMS1 | Hadha | Mus musculus | Trifunctional enzyme subunit alpha, mitochondrial | 20.5476211 | 20.4237365 | 0.123884647 |
| Q8BWT1 | Acaa2 | Mus musculus | 3-ketoacyl-CoA thiolase, mitochondrial | 21.6345554 | 21.512415 | 0.122140425 |
| P11440 | Cdk1 | Mus musculus | Cyclin-dependent kinase 1 | 20.4261476 | 20.3062417 | 0.119905877 |
| P20152 | Vim | Mus musculus | Vimentin | 23.4298194 | 23.3100802 | 0.119739244 |
| Q3UL36 | Arglu1 | Mus musculus | Arginine and glutamate-rich protein 1 | 22.0193428 | 21.9011483 | 0.118194494 |
| P62702 | Rps4x | Mus musculus | 40S ribosomal protein S4, X isoform | 22.5264547 | 22.4112284 | 0.115226269 |
| Q6ZWV7 | Rpl35 | Mus musculus | 60S ribosomal protein L35 | 21.4545694 | 21.3430077 | 0.111561673 |
| P62281 | Rps11 | Mus musculus | 40S ribosomal protein S11 | 22.9517674 | 22.8481793 | 0.103588171 |
| Q9Z2K1 | Krt16 | Mus musculus | Keratin, type I cytoskeletal 16 | 26.5897801 | 26.493811 | 0.095969059 |
| Q9JJI8 | Rpl38 | Mus musculus | 60S ribosomal protein L38 | 20.0944695 | 20.003214 | 0.091255484 |
| Q05816 | Fabp5 | Mus musculus | Fatty acid-binding protein 5 | 20.5347735 | 20.4459017 | 0.088871797 |
| P97351 | Rps3a | Mus musculus | 40S ribosomal protein S3a | 20.737627 | 20.6495993 | 0.088027635 |
| O35737 | Hnrnph1 | Mus musculus | Heterogeneous nuclear ribonucleoprotein H | 19.6356295 | 19.5530501 | 0.082579394 |
| P51410 | Rpl9 | Mus musculus | 60S ribosomal protein L9 | 22.9051542 | 22.8246982 | 0.080455982 |
| Q62186 | Ssr4 | Mus musculus | Translocon-associated protein subunit delta | 20.9554593 | 20.8829978 | 0.072461585 |
| Q60930 | Vdac2 | Mus musculus | Voltage-dependent anion-selective channel protein 2 | 20.1015972 | 20.0306467 | 0.070950522 |
| Q02566 | Myh6 | Mus musculus | Myosin-6 | 20.7177914 | 20.6472714 | 0.070519959 |
| Q9CY64 | Blvra | Mus musculus | Biliverdin reductase A | 20.0893413 | 20.0192548 | 0.070086579 |
| P02668 | CSN3 | Bos taurus | Kappa-casein | 22.3950762 | 22.3281784 | 0.066897783 |
| P13645 | KRT10 | Homo sapiens | Keratin, type I cytoskeletal 10 | 26.9976578 | 26.9315686 | 0.06608919 |
| P62908 | Rps3 | Mus musculus | 40S ribosomal protein S3 | 22.1139299 | 22.0503205 | 0.063609386 |
| P18760 | Cfl1 | Mus musculus | Cofilin-1 | 22.1022213 | 22.0397618 | 0.062459512 |
| Q02257 | Jup | Mus musculus | Junction plakoglobin | 20.2770331 | 20.2151486 | 0.061884518 |
| Q6IFZ6 | Krt77 | Mus musculus | Keratin, type II cytoskeletal 1b | 27.0914399 | 27.0296007 | 0.061839254 |
| O54734 | Ddost | Mus musculus | Dolichyl-diphosphooligosaccharide--protein glycosyltransferase 48 kDa subunit | 19.9614628 | 19.9014863 | 0.059976477 |
| Q8VDJ3 | Hdlbp | Mus musculus | Vigilin | 20.1343003 | 20.0766609 | 0.05763947 |
| P11679 | Krt8 | Mus musculus | Keratin, type II cytoskeletal 8 | 24.5638368 | 24.5105073 | 0.053329502 |
| P26443 | Glud1 | Mus musculus | Glutamate dehydrogenase 1, mitochondrial | 20.8805433 | 20.8279308 | 0.052612479 |
| P62751 | Rpl23a | Mus musculus | 60S ribosomal protein L23a | 20.5644434 | 20.5125599 | 0.051883497 |
| P11499 | Hsp90ab1 | Mus musculus | Heat shock protein HSP 90-beta | 21.8559701 | 21.804094 | 0.051876092 |
| Q61136 | Prpf4b | Mus musculus | Serine/threonine-protein kinase PRP4 homolog | 19.5598872 | 19.5112097 | 0.048677495 |
| P29699 | Ahsg | Mus musculus | Alpha-2-HS-glycoprotein | 19.5268348 | 19.4796704 | 0.047164497 |
| Q9CQM8 | Rpl21 | Mus musculus | 60S ribosomal protein L21 | 21.755212 | 21.7103486 | 0.044863407 |
| P61358 | Rpl27 | Mus musculus | 60S ribosomal protein L27 | 20.5426942 | 20.5010675 | 0.041626661 |
| P47754 | Capza2 | Mus musculus | F-actin-capping protein subunit alpha-2 | 20.4233206 | 20.3829191 | 0.040401539 |
| P09542 | Myl3 | Mus musculus | Myosin light chain 3 | 20.9371966 | 20.9029962 | 0.03420047 |
| Q91VR2 | Atp5f1c | Mus musculus | ATP synthase subunit gamma, mitochondrial | 21.5103784 | 21.4771788 | 0.033199647 |
| P27773 | Pdia3 | Mus musculus | Protein disulfide-isomerase A3 | 20.4482738 | 20.4174962 | 0.030777591 |
| Q6ZWU9 | Rps27 | Mus musculus | 40S ribosomal protein S27 | 21.5212301 | 21.5027937 | 0.018436365 |
| P51437 | Camp | Mus musculus | Cathelicidin antimicrobial peptide | 20.0358368 | 20.0179453 | 0.017891503 |
| E9PZF0 | Gm20390 | Mus musculus | Nucleoside diphosphate kinase | 20.9608507 | 20.9490045 | 0.011846179 |
| Q9CQ62 | Decr1 | Mus musculus | 2,4-dienoyl-CoA reductase [(3E)-enoyl-CoA-producing], mitochondrial | 19.5588777 | 19.5502051 | 0.008672577 |
| O08807 | Prdx4 | Mus musculus | Peroxiredoxin-4 | 22.8869517 | 22.8845917 | 0.002359958 |

Organoids

| **Protein ID** | **Gene** | **Organism** | **Description** | **V2GFP_log** | **WT_log** | **Subtraction value** |
| --- | --- | --- | --- | --- | --- | --- |
| D3YY75 | Vangl2 | Mus musculus | Vang-like protein | 21.84282 | 0 | 21.84282 |
| B1ARA5 | Rpl26 | Mus musculus | 60S ribosomal protein L26 | 17.9679 | 0 | 17.9679 |
| D3Z6P1 | Paics | Mus musculus | Phosphoribosylaminoimidazolesuccinocarboxamide synthase (Fragment) | 17.41143 | 0 | 17.41143 |
| E9Q1Y9 | Krt83 | Mus musculus | Keratin 83 | 17.32021 | 0 | 17.32021 |
| S4R2M7 | Pgk1 | Mus musculus | Phosphoglycerate kinase | 17.23216 | 0 | 17.23216 |
| A0A0N4SV32 | Serbp1 | Mus musculus | Plasminogen activator inhibitor 1 RNA-binding protein | 16.58524 | 0 | 16.58524 |
| A0A1D5RM76 | Tubb3 | Mus musculus | Tubulin beta-3 chain | 21.28915 | 19.3698266 | 1.919323 |
| A0A140LHK0 | Serpinh1 | Mus musculus | Serpin H1 (Fragment) | 19.94842 | 18.21034578 | 1.738079 |
| F2Z483 | Tcp1 | Mus musculus | T-complex protein 1 subunit alpha | 17.94602 | 16.35936394 | 1.586654 |
| J3QPZ9 | Eno3 | Mus musculus | Phosphopyruvate hydratase (Fragment) | 20.83612 | 19.31788732 | 1.518233 |
| E9Q6P8 | Rsrc1 | Mus musculus | Serine/Arginine-related protein 53 | 18.19932 | 16.90029919 | 1.299023 |
| Q542V3 | Srsf4 | Mus musculus | Serine/arginine-rich-splicing factor 4 | 19.55513 | 18.30309161 | 1.252039 |
| A0A0A0MQA5 | Tuba4a | Mus musculus | Tubulin alpha chain (Fragment) | 20.99879 | 19.8887431 | 1.110047 |
| A0A0G2JEY6 | Rpl34 | Mus musculus | 60S ribosomal protein L34 | 20.0776 | 18.97127278 | 1.106323 |
| A0A0A6YW33 | Gm6525 | Mus musculus | Predicted pseudogene 6525 | 19.96905 | 18.93774966 | 1.031298 |
| A0A0N4SUM2 | Hnrnpa2b1 | Mus musculus | Heterogeneous nuclear ribonucleoproteins A2/B1 (Fragment) | 18.27496 | 17.29576768 | 0.97919 |
| A0A2I3BQF4 | Rpl30 | Mus musculus | 60S ribosomal protein L30 | 18.10517 | 17.15349612 | 0.951674 |
| Q8BTU6 | Eif4a2 | Mus musculus | RNA helicase | 19.89662 | 18.98146226 | 0.915155 |
| A0A1L1SUK3 | Rpsa | Mus musculus | 40S ribosomal protein SA | 19.28043 | 18.55166814 | 0.728759 |
| A2BH06 | Rpl11 | Mus musculus | 60S ribosomal protein L11 (Fragment) | 21.38445 | 20.71263598 | 0.671815 |
| D3YWC2 | Arl6ip4 | Mus musculus | ADP-ribosylation factor-like protein 6-interacting protein 4 | 19.30507 | 18.64285494 | 0.662216 |
| D3Z2H9 | Tpm3-rs7 | Mus musculus | Tropomyosin 3, related sequence 7 | 18.77719 | 18.12881643 | 0.64837 |
| E9Q5F4 | Actb | Mus musculus | Actin, cytoplasmic 1 (Fragment) | 23.12012 | 22.47531591 | 0.644802 |
| A0A2R8VHH0 | Rac2 | Mus musculus | Ras-related C3 botulinum toxin substrate 2 (Fragment) | 17.9598 | 17.36543951 | 0.59436 |
| B1ATS4 | Atp2a3 | Mus musculus | Calcium-transporting ATPase | 15.40871 | 14.82797615 | 0.580733 |
| A0A2I3BPG9 | Rpl36a-ps1 | Mus musculus | Ribosomal protein L36A, pseudogene 1 | 23.16801 | 22.62038617 | 0.547626 |
| A0A1B0GRC0 | Col4a1 | Mus musculus | Collagen alpha-1(IV) chain (Fragment) | 17.67519 | 17.13679421 | 0.538394 |
| A0A0G2JEW4 | Prpf38b | Mus musculus | Pre-mRNA-splicing factor 38B | 19.05145 | 18.52473049 | 0.526715 |
| Q3UV11 | Krt6b | Mus musculus | Keratin, type II cytoskeletal 6B | 20.86416 | 20.34843581 | 0.515721 |
| E9Q509 | Pklr | Mus musculus | Pyruvate kinase | 19.2462 | 18.73320063 | 0.513002 |
| D3YZ68 | Eef1a1 | Mus musculus | Elongation factor 1-alpha 1 (Fragment) | 20.45617 | 19.96427518 | 0.491897 |
| J3QPS8 | Eif5a | Mus musculus | Eukaryotic translation initiation factor 5A-1 | 16.85956 | 16.37095196 | 0.488604 |
| A0A3B2WCD8 | Khsrp | Mus musculus | Far upstream element-binding protein 2 | 17.60561 | 17.13497122 | 0.470636 |
| A2AWQ2 | Rcc2 | Mus musculus | Protein RCC2 (Fragment) | 17.93335 | 17.46542973 | 0.467919 |
| F2Z456 | Cyb5r3 | Mus musculus | NADH-cytochrome b5 reductase | 20.21959 | 19.76018817 | 0.459399 |
| D3YYV8 | Rpl5 | Mus musculus | 60S ribosomal protein L5 (Fragment) | 18.92264 | 18.46326213 | 0.459375 |
| A0A2R8VHN4 | Rpl3 | Mus musculus | 60S ribosomal protein L3 (Fragment) | 21.15926 | 20.70109016 | 0.458168 |
| D3YVB4 | Rps15a | Mus musculus | 40S ribosomal protein S15a (Fragment) | 20.09508 | 19.63920397 | 0.455881 |
| A2A5N3 | Pabpc1l | Mus musculus | Polyadenylate-binding protein | 17.67165 | 17.2220169 | 0.449629 |
| A0A0G2JG40 | Gna12 | Mus musculus | Guanine nucleotide-binding protein subunit alpha-12 (Fragment) | 18.29641 | 17.85349079 | 0.442923 |
| B8JKK2 | Rpl15 | Mus musculus | Ribosomal protein L15 (Fragment) | 23.55659 | 23.11614035 | 0.440451 |
| A0A140T8T4 | Rpl9-ps6 | Mus musculus | 60S ribosomal protein L9 | 19.01937 | 18.60995145 | 0.409418 |
| B1ARR7 | Eno1 | Mus musculus | Alpha-enolase (Fragment) | 22.56958 | 22.18863351 | 0.380948 |
| A0A494B947 | U2af1 | Mus musculus | Splicing factor U2AF 35 kDa subunit (Fragment) | 18.95404 | 18.58012984 | 0.373907 |
| D3YV43 | Rps3 | Mus musculus | 40S ribosomal protein S3 | 20.72058 | 20.3530557 | 0.367526 |
| A0A494B9Z0 | Fau | Mus musculus | 40S ribosomal protein S30 | 23.59263 | 23.23246536 | 0.360169 |
| A0A1B0GSE8 | Rps11 | Mus musculus | 40S ribosomal protein S11 (Fragment) | 19.36066 | 19.01003391 | 0.350623 |
| E9Q715 | Luc7l2 | Mus musculus | Putative RNA-binding protein Luc7-like 2 | 19.65222 | 19.30729071 | 0.344934 |
| E9Q133 | Cct3 | Mus musculus | T-complex protein 1 subunit gamma | 17.95283 | 17.61756295 | 0.335271 |
| E9Q1F2 | Actb | Mus musculus | Actin, cytoplasmic 1 | 22.75222 | 22.43792601 | 0.314298 |
| H3BJ30 | Cpsf6 | Mus musculus | Cleavage and polyadenylation specificity factor subunit 6 | 19.23603 | 18.9441635 | 0.291863 |
| Q6ZWZ4 | Rpl36 | Mus musculus | 60S ribosomal protein L36 | 19.77603 | 19.49788953 | 0.278139 |
| A0A2R8W6V7 | Myh9 | Mus musculus | Myosin-9 (Fragment) | 19.80869 | 19.54550879 | 0.263179 |
| A0A494B945 | Rab1b | Mus musculus | Ras-related protein Rab-1B (Fragment) | 19.36052 | 19.10214842 | 0.25837 |
| Q5SQG5 | Phb | Mus musculus | Prohibitin (Fragment) | 17.99661 | 17.74284539 | 0.25377 |
| Q14AA6 | 1700009N14Rik | Mus musculus | GTP-binding nuclear protein Ran | 22.49095 | 22.24171605 | 0.249231 |
| Q504P4 | Hspa8 | Mus musculus | Heat shock cognate 71 kDa protein | 20.96138 | 20.71446057 | 0.246923 |
| E9Q3D6 | Hsp90ab1 | Mus musculus | Heat shock protein HSP 90-beta (Fragment) | 18.55704 | 18.3487314 | 0.208311 |
| F6XI62 | Rpl7 | Mus musculus | 60S ribosomal protein L7 (Fragment) | 26.90885 | 26.71292828 | 0.19592 |
| Q6PHC1 | Eno1 | Mus musculus | Phosphopyruvate hydratase | 20.92055 | 20.72555041 | 0.194996 |
| Q3UH59 | Myh10 | Mus musculus | Myosin-10 | 19.84983 | 19.65520094 | 0.194627 |
| I7HLV2 | Rpl10 | Mus musculus | 60S ribosomal protein L10 (Fragment) | 21.28471 | 21.09105206 | 0.193654 |
| A0A1D5RLW5 | Rpl18a | Mus musculus | 60S ribosomal protein L18a | 21.67333 | 21.49239505 | 0.180934 |
| D3Z6F5 | Atp5a1 | Mus musculus | ATP synthase subunit alpha | 21.24223 | 21.09752851 | 0.144699 |
| A0A087WNS0 | Rpl3 | Mus musculus | 60S ribosomal protein L3 | 21.52243 | 21.38351032 | 0.138919 |
| A2A6F8 | Rpl23 | Mus musculus | 60S ribosomal protein L23 (Fragment) | 20.21196 | 20.07419769 | 0.137758 |
| E9QN70 | Lamb1 | Mus musculus | Laminin subunit beta-1 | 21.76032 | 21.63409454 | 0.126229 |
| A2A547 | Rpl19 | Mus musculus | Ribosomal protein L19 | 24.83364 | 24.71030281 | 0.123339 |
| A0A1B0GS68 | Gm45713 | Mus musculus | 60S ribosomal protein L13a | 23.1214 | 23.00672726 | 0.11467 |
| B0V2N7 | Anxa2 | Mus musculus | Annexin (Fragment) | 19.51767 | 19.40895654 | 0.10871 |
| A0A1L1SU37 | Pkm | Mus musculus | Pyruvate kinase (Fragment) | 20.50187 | 20.39582195 | 0.106047 |
| B1B0C7 | Hspg2 | Mus musculus | Basement membrane-specific heparan sulfate proteoglycan core protein | 21.06258 | 20.95963347 | 0.102948 |
| A0A1L1SRW0 | Rpsa | Mus musculus | 40S ribosomal protein SA (Fragment) | 19.28661 | 19.18552331 | 0.101082 |
| A0A3Q4L393 | Srsf7 | Mus musculus | Serine/arginine-rich-splicing factor 7 | 19.68363 | 19.6001082 | 0.083519 |
| P35527 | KRT9 | Homo sapiens | Keratin, type I cytoskeletal 9 | 25.23352 | 25.15267229 | 0.08085 |
| F6ZSB7 | Phgdh | Mus musculus | D-3-phosphoglycerate dehydrogenase (Fragment) | 17.85064 | 17.78112191 | 0.069522 |
| B2M1R6 | Hnrnpk | Mus musculus | Heterogeneous nuclear ribonucleoprotein K | 17.75568 | 17.69214817 | 0.063534 |
| A0A1B0GSB2 | Rpl13a | Mus musculus | 60S ribosomal protein L13a | 22.31871 | 22.26082121 | 0.057888 |
| A0A0G2JDW7 | Rps27 | Mus musculus | 40S ribosomal protein S27 (Fragment) | 19.27528 | 19.22828258 | 0.046997 |
| Q921R2 | Rps13 | Mus musculus | 40S ribosomal protein S13 | 18.71505 | 18.69286768 | 0.022185 |
| Q9D8L3 | Ssr4 | Mus musculus | Translocon-associated protein subunit delta | 16.78551 | 16.76751692 | 0.017989 |
| D3Z2F2 | Hspd1 | Mus musculus | 60 kDa heat shock protein, mitochondrial (Fragment) | 18.42755 | 18.4122763 | 0.015275 |
| E9PUU2 | Clu | Mus musculus | Clusterin (Fragment) | 19.6523 | 19.64199509 | 0.010302 |
